# Deep mapping of structural perturbations to energetics enables precise TCR design

**DOI:** 10.64898/2026.04.15.718814

**Authors:** Shifu Luo, Songming Zhang, Ying Shi, Junjie Li, Jiaxin Cai, Ningyi Shao, Yi Pan, Jinyan Li

## Abstract

Affinity optimization and cross-reactivity profiling are pivotal for T-cell receptor (TCR) engineering and immunotherapy, yet remain hindered by the vast diversity of the TCR repertoire and limited structural insights. Here, we present mpTCRai, a deep learning framework that captures the sequential characteristics of antigen presentation and recognition with high precision. Leveraging these precise structural predictions, we established a contact hotspot-based scoring mechanism that explicitly maps structural perturbations to energetic changes, yielding a strong correlation with experimental affinity (r = −0.88). In application, the model identified critical molecular switches, such as specific G-to-A substitutions, and revealed their dependence on structural context. Furthermore, it effectively distinguished functional variants from dangerous cross-reactive mutations like Y5W, thereby mitigating the risk of off-target toxicity. Guided by these insights, we computationally designed four novel A6-TCR variants, demonstrating a rational strategy for candidate selection in adult T-cell leukemia. This work establishes a unified platform integrating structural and energetic constraints to advance precise, safe TCR therapeutic design.

## Introduction

The activation of CD8+ T cells plays decisive roles in antigen-specific immune responses against infected or malignant cells^1–3^. This process goes through two sequential stages: (i) antigen presentation as mediated by major histocompatibility complex class I (MHC-I) molecules for loading the peptide antigens to the cell surface, and (ii) T cell recognition as driven by T cell receptor (TCR) binding to the peptide–MHC (pMHC) complexes^4^. Decoding the specificity of TCR–pMHC interactions is critical for advancing TCR-based immunotherapies. For instance, engineering high-affinity TCRs has emerged as a crucial strategy for enhancing T cell-mediated antitumor responses, as binding affinity directly correlates with therapeutic efficacy^5,6^.

However, TCRs possess a remarkable diversity due to combinatorial V(D)J recombination and junctional diversity, creating a theoretical repertoire exceeding 10^1^□ unique variants^7–9^. Current characterization of TCR-pMHC interactions remains constrained by low-throughput structural and functional assays^10^, which are inadequate for systematic exploration of this vast sequence space. These limitations hinder a mechanistic understanding of TCR recognition and pose huge challenges for the development of TCR-based immunotherapies.

Conventional approaches typically employ affinity maturation and structure-guided design to optimize TCR properties, investigate biological mechanisms, and develop clinical candidates^11^. Yet each step of this process faces substantial difficulties: affinity optimization requires costly and time-consuming experimental screening to identify functionally relevant mutations, while safety assessment necessitates comprehensive cross-reactivity profiling across the vast peptidome spaces^12,13^. Thus, reliable predictive tools for assessing mutation effects and off-target potentials are highly demanded in developing safe, effective TCR-based therapeutics.

Existing computational tools in this domain can be broadly categorized into two methodological paradigms based on their predictive outputs. The first category comprises sequence-based classification models. These tools address distinct biological tasks, including the prediction of precise TCR-epitope binding specificity (e.g., NetTCR^14^, epiTCR^15^) and the assessment of broader peptide immunogenicity or neoantigen presentation (e.g., PRIME^16^, DeepNeo-TCR^17^). Despite these different objectives, they share a common computational approach. They rely exclusively on 1D sequence information from human leukocyte antigen (HLA) molecules, antigens, and TCRs to perform a binary classification (e.g., binding vs. non-binding, or immunogenic vs. non-immunogenic). The second category employs regression-based methods to predict residue-level distances between TCRs and antigens, thereby providing explicit structural information with direct relevance to rational TCR engineering and vaccine design.

The recent TEIM^18^ method is an advanced tool in the area of structure-based TCR specificity prediction, yet its accuracy is compromised by the narrow availability of structural data and inherent constraints in its modeling architecture. Specifically, the model’s framework does not consider the sequential nature of the T cell activation process; it just focuses on direct TCR-antigen interactions while neglecting the fundamental antigen-presentation principle. These shortcomings underscore critical need for more biology-motivated predictive algorithms and needs for expanded high-quality datasets to drive significant progress in TCR engineering and computational immunology.

Here, we developed a novel deep learning framework—mpTCRai (mutual-perspective TCR–antigen interaction model)—for accurate TCR-antigen specificity prediction. The framework explicitly models the sequential, two-stage process of T cell activation through dual-representation learning and incorporates a unique mutual-perspective cross-attention (MPCA) mechanism to capture the bidirectional recognition dynamics between TCR and pHLA (**Fig. 1a**). Specifically, during the antigen presentation stage, mpTCRai employs ESM-2^19^ to extract HLA and antigen sequence representation vectors and then utilizes an HLA-peptide interaction module to simulate the antigen binding and presentation process, thereby generating a comprehensive feature vector of the pHLA complex (denote it as **P**). For modeling the T cell recognition stage, the framework employ TCR-BERT^20^ to encode TCR CDR3β sequences (denote it as **T**), followed by a TCR-pHLA interaction module that alternately captures recognition patterns from both the TCR-centric and pHLA-centric viewpoints. The mutual interaction is featured to compute as: ℱ _interaction_ *f* (*Attention*_TCR_view_ (*Q* = ***T***, *K* = ***P***, *V* = ***P***)) ⊙ *f* (*Attention*_pHLA_view_(*Q* = ***P***, *K* = ***T***, *V =* ***T*** (see details in the **Methods** section).

**Figure 1.**
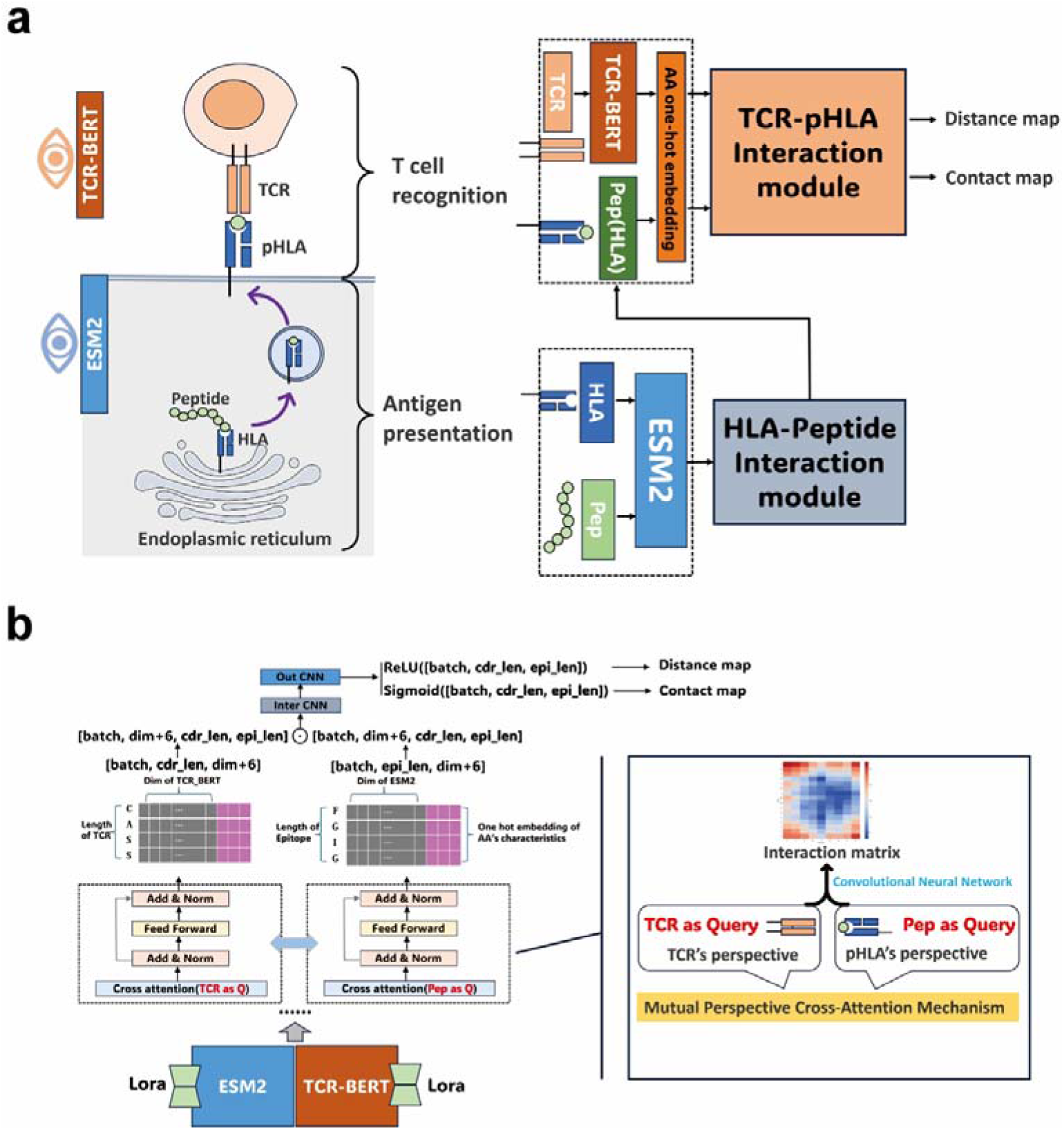
Architecture of mpTCRai for modeling and prediction of TCR-antigen interaction. **a**. Biologically inspired two-stage workflow. The model mimics the natural immunological process by sequentially simulating antigen presentation (pHLA) and T cell recognition. Pre-trained encoders (ESM2 for pHLA, TCR-BERT for TCR) generate initial embeddings, which are processed through specialized interaction modules to capture domain-specific features. **b**. Architecture of the Mutual-Perspective Cross-Attention module. This core mechanism establishes a bidirectional information flow, iteratively alternating between TCR-centric and pHLA-centric attention views. This design allows the model to capture fine-grained, reciprocal dependencies at the residue level, effectively aligning sequence features with structural contexts.

This MPCA mechanism alternating between the TCR-centric and pHLA-centric views can effectively capture complementary interaction patterns (**Fig. 1b, S1**). These interaction features are then processed through a Resnet^21^ module to generate residue-distance matrices and contact-probability maps.

To alleviate the bottleneck of data scarcity and establish a reliable benchmark for model validation, we curated a comprehensive dataset of structurally characterized TCR–peptide–MHC ternary complexes. Evaluated on this dataset, mpTCRai demonstrated robust predictive performance while maintaining biological interpretability by explicitly modeling the sequential stages of the immunological activation process. Through two detailed case studies, we showed that mpTCRai can identify key molecular switches that modulate binding affinity and flag potential cross-reactive mutants, offering actionable insights for TCR engineering and vaccine safety assessment. To demonstrate the translational utility of our framework, we focused on Adult T-cell leukemia (ATL), an aggressive hematologic malignancy caused by HTLV-1 infection where the Tax peptide serves as a key immunotherapeutic target^22^. By applying mpTCRai to perform combinatorial mutagenesis on the A6-TCR, we computationally engineered four TCR variants with enhanced binding affinity, providing a novel rational design strategy to accelerate the development of ATL therapeutics.

Overall, the framework establishes robust connections between sequence information, structural features, and binding energetics, offering an effective computation platform for developing and optimizing TCR-based therapeutics.

## Results

### Prediction Performance by mpTCRai

Given the few-shot learning^23^ nature of our task, we fine-tuned mpTCRai directly on the STCRDab dataset^24^ (n=90). Specifically, all pre-trained parameters within the ESM-2 and TCR-BERT representation layers were frozen and fine-tuned exclusively using Low-Rank Adaptation (LoRA^25^), while the parameters within the interaction layer remained fully trainable. We subsequently evaluated the trained model on two independent test sets: the test1 (n=7) dataset and a newly constructed test2 (n=67) dataset (see details at the **Methods** section). Although our training set contains only 90 samples, it is important to emphasize that these samples are a subset of the original TEIM training data, thereby ensuring a fair model comparability (**Fig. 2a**).

**Figure 2.**
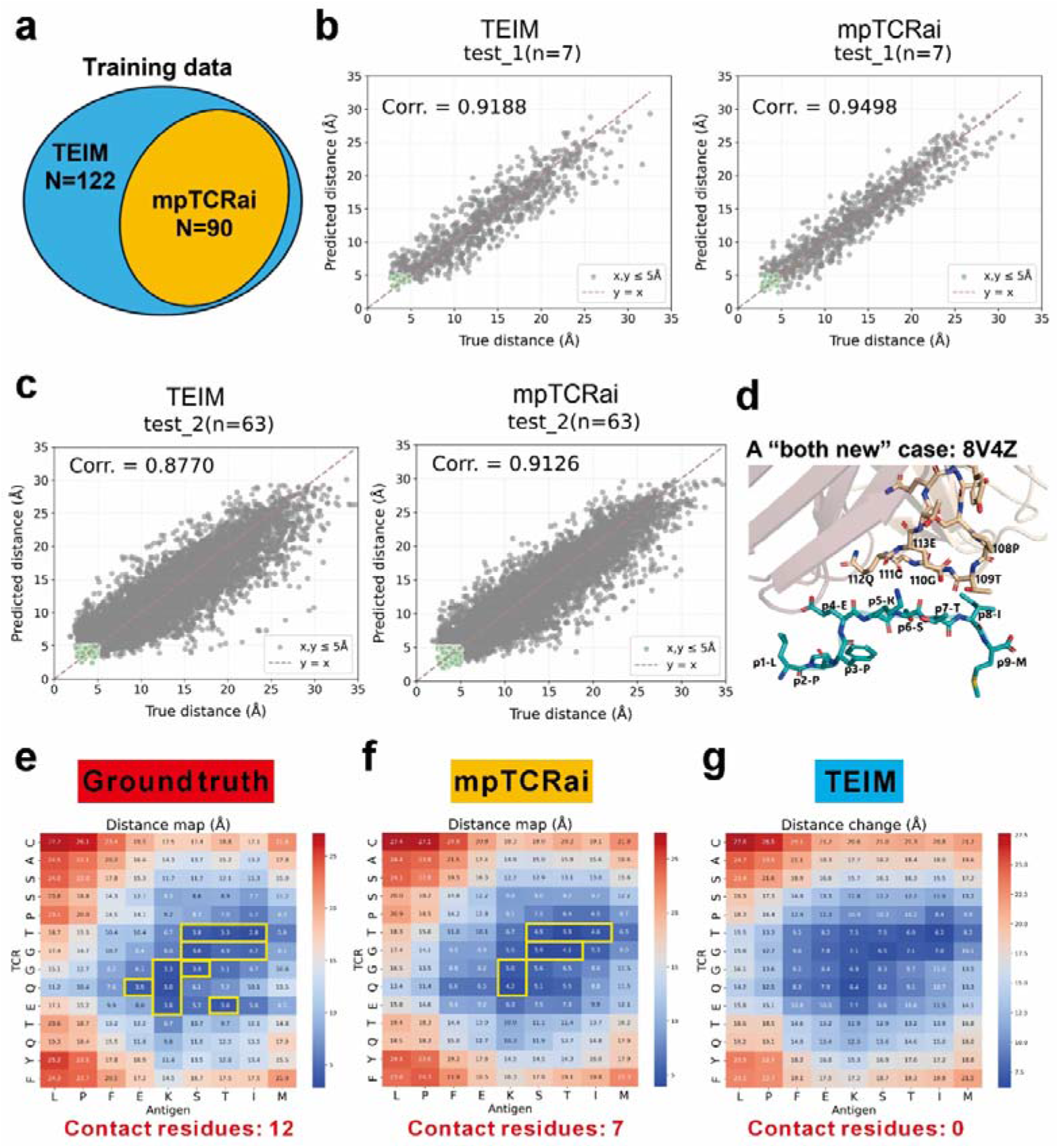
Superior structural modeling and generalization capability of mpTCRai. **a**. Comparison of fine-tuning data scale between TEIM and mpTCRai. **b**. Correlation analysis between ground-truth and predicted residue distances on the test1 dataset. **c**. Correlation analysis between ground-truth and predicted residue distances on the test2 dataset. **d**. 3D structural visualization of the unseen TCR-antigen complex (PDB: 8V4Z). **e**. Ground truth contact map highlighting the 12 experimental contact residues (yellow box). **f**. mpTCRai prediction heatmap, identifying seven verified true contacts (yellow box). **g**. TEIM prediction heatmap, showing a failure to capture any valid contact residues.

The mpTCRai takes the sequences of the HLA–antigen–TCR ternary complex as input and outputs a residue distance matrix between the TCR and antigen to represent their interaction landscape. On the test1 dataset, mpTCRai has achieved superior performance in comparison with TEIM, having a Pearson correlation coefficient of 0.9498 between the predicted and actual residue-pair distances (versus TEIM’s 0.9188; **Fig. 2b**). In the interaction prediction for contact residues of distance ≤5□□, mpTCRai made a 15 percentage-point improvement in recall while maintaining comparable precision (**Table 1**).

**Table 1.** Prediction performance of mpTCRai on two test datasets in comparison with TEIM.

| Dataset | Model | Accuracy (%) | Precision (%) | Recall (%) | F1-score (%) |
| --- | --- | --- | --- | --- | --- |
| **Test1**<br>(n=7) | TEIM | 93.47 | **72.09** | 38.75 | 50.22 |
|  | mpTCRai | **94.11** | 70.49 | **53.75** | **60.98** |
| | $\Delta$ | **+0.64** ↑ | **-1.60** ↓ | **+15.00** ↑ | **+10.76** ↑ |
| **Test2**<br>(n=67) | TEIM | 93.00 | 57.98 | 21.13 | 31.02% |
|  | mpTCRai | **93.62** | **61.37** | **38.44** | **47.26%** |
| | $\Delta$ | **+0.62** ↑ | **+3.39** ↑ | **+17.31** ↑ | **+16.25** ↑ |

To further assess robustness, we conducted evaluation on test2. TEIM attained a Pearson correlation of about 0.88 between the predicted and actual residue-pair distances, consistent with its original reported performance^18^, whereas mpTCRai achieved a higher correlation of 0.91 (**Fig. 2c**), preserving its >3% performance advantage as observed in test1. Importantly, for contact residue prediction, mpTCRai demonstrated a substantial 17.31-percentage-point gain in recall while simultaneously improving accuracy, precision, and F1-score (**Table 1**). These consistent results across distinct and independent test sets underscore mpTCRai’s state-of-the-art predictive performance despite limited training data, highlighting the superiority of mutual perspective framework.

We use a detailed example to demonstrate that mpTCRai’s generalization capability is much better than TEIM. This example is a “both new” case (PDB ID: 8V4Z^26^) whose antigen and TCR sequences are both absent from the training data. In fact, its native structure has 12 contact residues compacted between the TCR positions from 109T-112Q and the antigen positions from 5K-8I (**Fig. 2d&e**). While TEIM cannot predict any correct contacts (predicted contacts = 0; **Fig. 2g**), mpTCRai successfully identified seven contact pairs that largely overlap with the core interaction interface (**Fig. 2f**). This example illustrates our model’s capacity to generalize to novel sequences and identify their key interaction hotspots.

### Linking Residue Mutations to Energetic Changes (ΔΔG) through Structural Perturbation

To gain deeper insights into TCR-antigen interactions, we applied mpTCRai to analyze a series of CDR3β and epitope variants. We investigated how residue mutations cause structural perturbations, how these conformational changes affect binding energy, and sought to bridge structural alterations to energetic changes.

Our study is on the well-characterized A6 TCR (CASRPGLAGGRPEQYF) in complex with the Tax peptide (LLFGYPVYV), the first solved human TCR-pHLA structure^27–29^. The experimental ΔΔG measurements of its mutants are public available and the same as A6-Tax mutation dataset^18^ which has been previously used by TEIM for performance evaluation (**Supplementary Table 4**).

For every mutant in this dataset, we used mpTCRai (with the HLA, antigen, and TCR sequences as input data) to generate the corresponding TCR–antigen residue-pair distance matrix. After computing the difference between the distance matrix of the mutant and that of the reference (wild-type), we got a residue-level distance change matrix to quantify the mutation-induced structural perturbations. In this matrix, positive values indicate residue-pair separation (weakened interactions), whereas negative values stand for inter-residue proximity (enhanced interactions). The overall structural perturbation value for each mutant is calculated by summing all elements in the distance change matrix and then taking the mean of the resulting sum. Then, we correlated the overall structural perturbation values of all the mutants with their ΔΔG values. Our analysis revealed a strong correlation (r = 0.70, **Fig. 3a**) between the predicted residue distance changes and the experimental binding affinity changes (ΔΔG); this correlation surpasses TEIM’s performance (a correlation of 0.65).

**Figure 3.**
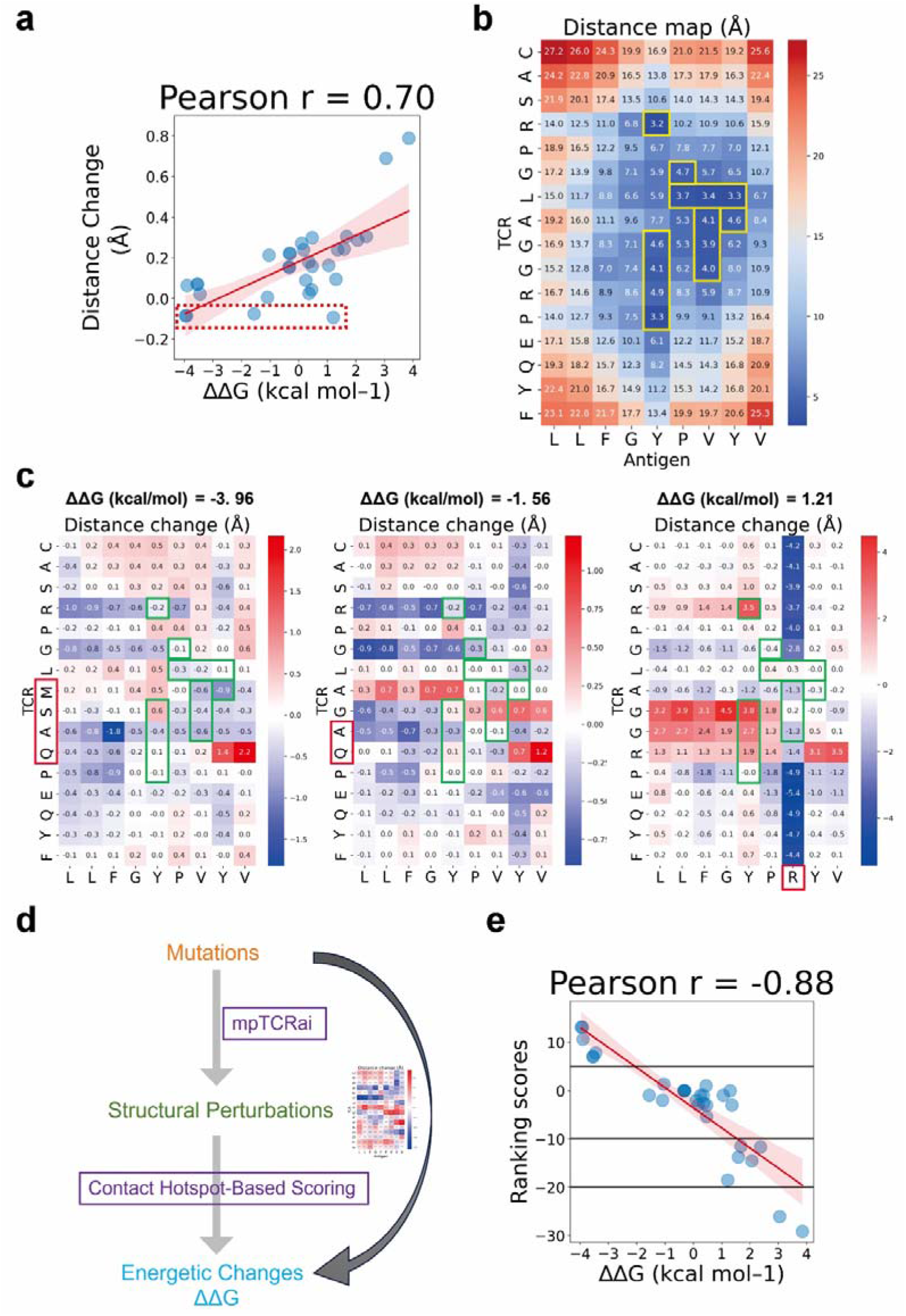
Structural perturbation modeling bridges sequence mutations to binding energetic landscapes. **a**. Quantitative relationship between predicted residue distance changes and experimental binding affinity changes (ΔΔG) for 31 A6-Tax mutants. **b**. Delineation of contact hotspots, defined as residue pairs within a 5 Å threshold, mapped onto the A6-Tax distance matrix (highlighted in yellow boxes). **c**. Residue distance change heatmaps for three representative mutants (indicated by red box in *a*), each showing distinct perturbation patterns within contact hotspot regions despite similar global distance changes. **d**. A schematic illustration of the link between mutations and energetic changes (ΔΔG). Showing how structural perturbations bridge sequence mutations with binding affinity. **e**. Validation of the contact hotspot-based scoring function, demonstrating a strong correlation with experimental ΔΔG values, thereby confirming the predictive power of structural perturbation mapping. Heatmap interpretation in *c*: blue indicates distance reduction (potential interaction strengthening); red indicates distance increase (potential interaction weakening).

Although mpTCRai has made the above overall robust performance, accurate prediction remains challenging for some mutants, particularly for those as indicated by the red box in **Fig. 3a**. As illustrated in **Fig. 3c**, three representative cases exhibit similar global distance changes but have markedly different ΔΔG values, ranging from –3.96 to +1.21, a critical phenomenon that has been overlooked in previous studies.

To dissect this discrepancy which may have been caused by the distance difference computed for all the residue pairs, we narrowed down to concentrate on only those contact residues (hotspot) which are defined as residue pairs within 5 Å in the wild-type complex (indicated by the yellow box in **Fig. 3b**). With this residue scope narrowing down, we found distinct perturbation patterns within these hotspot regions: Case 1 exhibited partial stabilization with closer proximity in specific hotspots (ΔΔG = –3.96); Case 2 maintained stability across most hotspots (ΔΔG = –1.56); and Case 3 displayed widespread disruption of hotspot interactions (ΔΔG = +1.21). Importantly, these structural alterations within hotspot regions align consistently with the observed energetic changes, indicating that local interaction variations in contact areas may serve as the primary driver of affinity modulation.

We further extended the analysis to cover additional representative cases and have found distinct structural patterns that underline different mutational effects. For example, mutations causing substantial affinity loss (ΔΔG > 2) consistently disrupted contact hotspots, exhibiting significantly increased residue-pair distances indicative of interaction breakdown (**Fig. S2**).

Based on these observations, we formulated a contact hotspot-based scoring function (**Methods**) that correlates strongly with experimental binding affinity changes (Pearson r = −0.88 between predicted scores and ΔΔG values) and it thus enables an effective stratification of functional variants (**Fig. 3e**). This approach establishes a bridge connecting sequence mutations to structural perturbations and ultimately to energetic changes, providing an integrated framework for interpreting TCR–antigen interactions (**Fig. 3d**).

### Identification of Affinity-enhancing Molecular Switches for A6-TCR Engineering

Conventional TCR optimization strategies, such as random mutagenesis and sequence-based analysis, are often inefficient and costly due to the lack of a mechanism-driven rationale for identifying key molecular switches that modulate binding affinity^30,31^. As illustrated in Figure 4a, obtaining a high-activity A6-TCR variant requires iterative rounds of sequence engineering and functional screening, with the mutated amino acid residues highlighted in red. Specifically, incorporating an additional ‘A’ mutation to transition from the “MS” mutant to the “MSA” mutant induces an abrupt and substantial enhancement in binding affinity toward the Tax epitope. However, sequence information alone cannot elucidate the mutagenesis underlying this affinity improvement.

**Figure 4.**
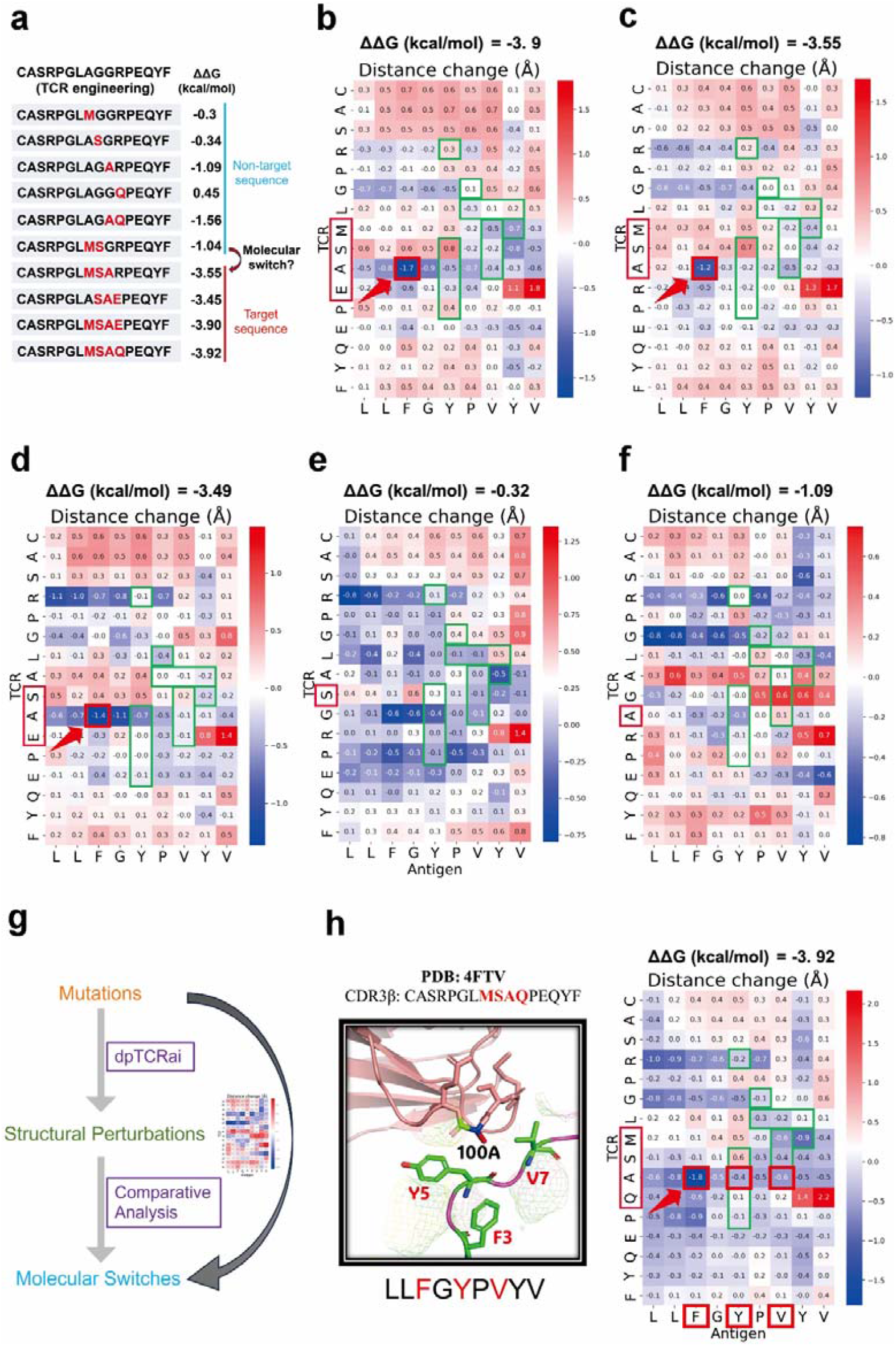
Structural and energetic determinants of TCR binding affinity as revealed through mpTCRai-guided engineering analysis. **a**. Mutated sequences and corresponding binding energy changes (ΔΔG) for A6-TCR variants. The Tax epitope remains unchanged across all the variants. **b-f**. Predicted residue distance change matrices for A6-TCR variants: (**b**) 98-101MSAE, (**c**) 98-100MSA, (**d**) 99-101SAE, (**e**) 99S, (**f**) 100A. Target variants (**b–d**) share similar distance change features at position 100A (highlighted with red arrow). **g**. Schematic illustration of the link between mutations and molecular switches, showing how structural perturbations enable the identification of molecular switches. **h**. Structural validation against the high-affinity A6-c134 mutant. (**Left**) Crystal structure (PDB: 4FTV) revealing the hydrophobic pocket formed by Tax residues 3F, 5Y, and 7V that spatially accommodates TCR residue 100A. (**Right**) The mpTCRai-predicted perturbation matrix for A6-c134 mirrors the pattern observed in target variants (\***b-d**\*), accurately capturing the significant proximity (blue region) between TCR 100A and the hydrophobic pocket. Heatmap interpretation: blue indicates reduced distances (potential interaction strengthening); red indicates increased distances (potential interaction weakening).

Our approach establishes a direct link between sequence mutations and residue-level structural perturbations, as described in the previous subsection: mpTCRai takes mutated TCR sequences along with the paired antigen and HLA sequences as input and outputs residue distance matrices. Mutation-induced structural perturbations are then quantified by subtracting the wild-type matrices from these mutant matrices. These structural perturbations constitute the structural basis of mutational effects (**Fig. 4g**).

We found that the GG-to-SA mutation at positions 99-100 is conserved across all the target TCR engineering sequences (**Fig. 4a**). To understand more about this conservation, we conducted analysis on the residue distance change matrices across all the target sequences. It was revealed that although the contact hotspot regions (highlighted in green box) remain largely unchanged, a consistent and significant reduction in inter-residue distance (1.2–1.7 Å) occurs specifically between TCR position 100 and Tax residue 3F. This spatial contraction likely facilitates new hydrophobic interactions between alanine and phenylalanine (A–F pair, red arrow) for enhancing binding affinity (**Fig. 4b-d**). In contrast, single point mutations (99S or 100A), while preserving the overall architecture of the contact hotspot regions similarly as the functional variants do, are unable to reconstitute the critical distance reduction between TCR position 100 and Tax 3F as observed in the functional variants (**Fig. 4e&f**). This inability to induce the key structural perturbation explains their limited effect on affinity. These findings suggest that the 100G-to-A substitution—not the 99G-to-S mutation—acts as a conserved molecular switch for the affinity modulation with its functional impact dependent on the cooperative interactions around the neighboring residues.

We further substantiated this finding using the crystal structure of the A6-c134 TCR–Tax complex (PDB: 4FTV; CDR3β: CASRPGL**MSAQ**PEQYF). Structural analysis revealed that Tax residues Phe3, Tyr5, and Val7 form a hydrophobic FYV pocket that accommodates the methyl group of Ala100 in the TCR’s CDR3β loop to stabilize a compact hydrophobic core (**Fig. 4g**). Predictive results from mpTCRai align closely with these structural observations. The residue distance change matrix generated by mpTCRai exhibits marked spatial proximity between Ala100 in A6-c134 TCR and the constituent residues of the Tax-3F5Y7V hydrophobic pocket (**Fig. 4h**). The concordance between these computational predictions and the experimental structural data underscores the functional importance of this hydrophobic interface in reinforcing the TCR binding. Notably, the predicted distance change matrices consistently capture this conserved structural motif across all the target sequences (**Fig. 4b–d**), providing a unified, structurally grounded explanation for the observed affinity enhancement. As further evidence, the deep mutational scanning data from Daniel T et al.^32^ have also confirmed that the A6-c134 TCR G-to-A substitution at position 100 is both necessary and sufficient to enhance TCR activity.

Collectively, these results highlight the robust potential of mpTCRai for TCR binding affinity optimization. Our approach not only supports accurate stratification of mutations and precise identification of functional TCR variants but also uncovers critical molecular switches and energetically favorable interaction mechanisms, offering actionable insights for rational TCR engineering.

### Cross-reactivity Profiling by mpTCRai

Cross-reactivity evaluation is an essential step prior to the clinical application of neoantigen vaccines or TCR-T therapies^33,34^. Accurate prediction of potential cross-reactivity with off-target cells or tissues is important otherwise the TCR mutants may result in severe, even fatal, adverse events^35–38^.

Previous structural and biochemical evidence^27,39^ indicates that the A6-TCR binding interface contains a specialized deep pocket that preferentially accommodates aromatic residues at the Tax peptide position Y5 (**Fig. 5b**). Functional profiling corroborates this structural preference, showing markedly reduced activity when non-aromatic residues occupy this position (**Fig. S3a**). Therefore, we classified substitutions of Tax-Y5 into two functional categories: aromatic-preserving “small-effect” variants, which maintain native-like cross-reactivity, and non-aromatic “large-effect” variants, which exhibit substantially diminished binding and specificity (**Fig. 5a**).

**Figure 5.**
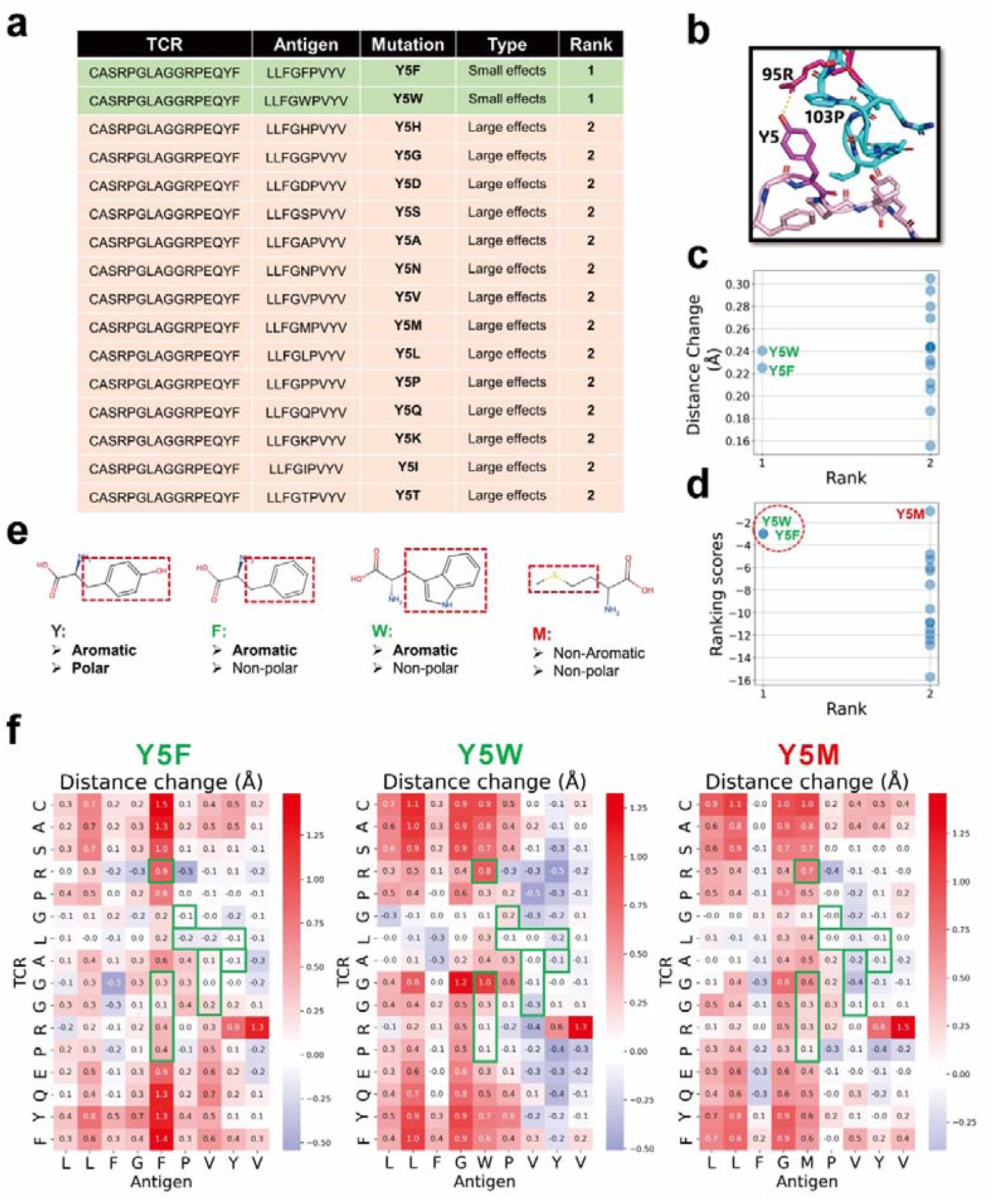
Dissecting the structural basis of TCR specificity and discriminating functional variants at the critical Tax Y5 position. **a**. Phenotypic categorization. Classification of Tax Y5 substitutions into activity-preserving (“small effect”) and disruptive (“large effect”) groups based on experimental activity. **b**. Structural anchor of specificity. Crystal structure of the A6-Tax complex (PDB: 1AO7) reveals the critical hydrogen bond between the Tax Y5 hydroxyl group and TCR residue 95R, establishing a key recognition interface. **c**. Insufficiency of global metrics. Scatter plot demonstrating that aggregate global distance changes fail to distinguish functional variants from disruptive ones. **d**. Effective stratification by hotspot scoring. The contact hotspot-based scoring function successfully separates mutation classes, correctly prioritizing functional variants like Y5M, Y5F, and Y5W as top-ranked candidates. **e**. Physicochemical compatibility. Analysis of top-ranking variants highlights the preservation of key properties (e.g., aromaticity) required for binding. **f**. Conserved structural signatures. Heatmaps of high-scoring variants reveal consistent perturbation patterns within contact hotspots (green boxes), indicating that these mutations maintain the essential interface geometry despite local variations.

Based on comprehensive amino acid substitution data of Tax-Y5, we conducted cross-reactivity assessments to validate our model and hotspot-based scoring function. Our objective was to recognize aromatic variants Y5F and Y5W among all the mutants, which have been previously documented to exhibit cross-reactivity with the Tax epitope. The analysis process consisted of two stages: first, mpTCRai processed the mutated sequences to generate residue distance change matrices that quantify mutation-induced structural perturbations; second, the hotspot-based scoring function utilized these matrices to score and rank all the mutants.

The results show that we successfully identified Y5F and Y5W as top-rank mutants, with ranking scores slightly below that of the Tax peptide (ranking score = 0) (**Fig. 5d**). In contrast, conventional analysis using global distance metrics failed to distinguish between “small-effect” and “large-effect” variants (**Fig. 5c**). These findings support two fundamental biological principles: first, that Y5 represents an evolutionary optimum, as all the substituted variants scored lower than the native residue; and second, that position 5 of the Tax epitope exhibits a strong preference for aromatic residues.

However, the model misclassified Y5M as being cross-reactive. This motivated us to take a deeper investigation into this discrepancy. Structural analysis revealed that the native Y5 achieves optimal recognition through its dual physicochemical properties—aromaticity and polarity—with its hydroxyl group forming an essential hydrogen bond with R95 on the A6 TCR (**Fig. 5b**). Although Y5F and Y5W retain aromatic character, their nonpolar side chains and altered conformations disrupt this key hydrogen bond, resulting in moderately reduced activity. In contrast, Y5M induces minimal structural perturbation but lacks both aromaticity and polarity, accounting for its functional impairment (**Fig. 5e&f**). This illustrates that the absence of essential physicochemical features can lead to significant functional deficits, even when overall structural distances appear favorable.

This case study further validates the robustness of our methodology. By establishing a predictive framework that connects sequence mutations to structural perturbations and to energetic consequences, we can effectively identify mutations with cross-reactive potential, thereby mitigating the risk of adverse clinical events. However, we note that our structural analysis represents only the initial screening. A complete pipeline for evaluating true off-target toxicity must involve scanning these mpTCRai-predicted candidates against the human proteome, followed by rigorous experimental validation, to ensure clinical safety.

### mpTCRai-empowered Large Scale Mutagenesis Study on A6-TCR

Based on our previously identified key molecular regulatory mechanisms—including the hydrophobic core interaction between TCR residue 100A and the Tax peptide’s hydrophobic pocket, and the critical hydrogen bond between TCR residue 94R and the Y5 residue of the Tax peptide—we established a structure-informed framework for evaluating the functional activity of TCR variants.

We conducted combinatorial mutagenesis experiments on the A6-TCR. In the first set, we introduced mutations at positions 8, 9, and 10 (corresponding to residues 98, 99, and 100 in the full-length sequence), generating a library of 7,999 mutant TCRs. In the second set, we targeted on positions 9, 10, and 11 (corresponding to residues 99, 100, and 101), constructing another library of 7,999 mutants (**Fig. 6a**). This multi-site, multi-combination parallel approach enables a comprehensive exploration of synergistic effects of mutations on TCR affinity and specificity.

**Figure 6.**
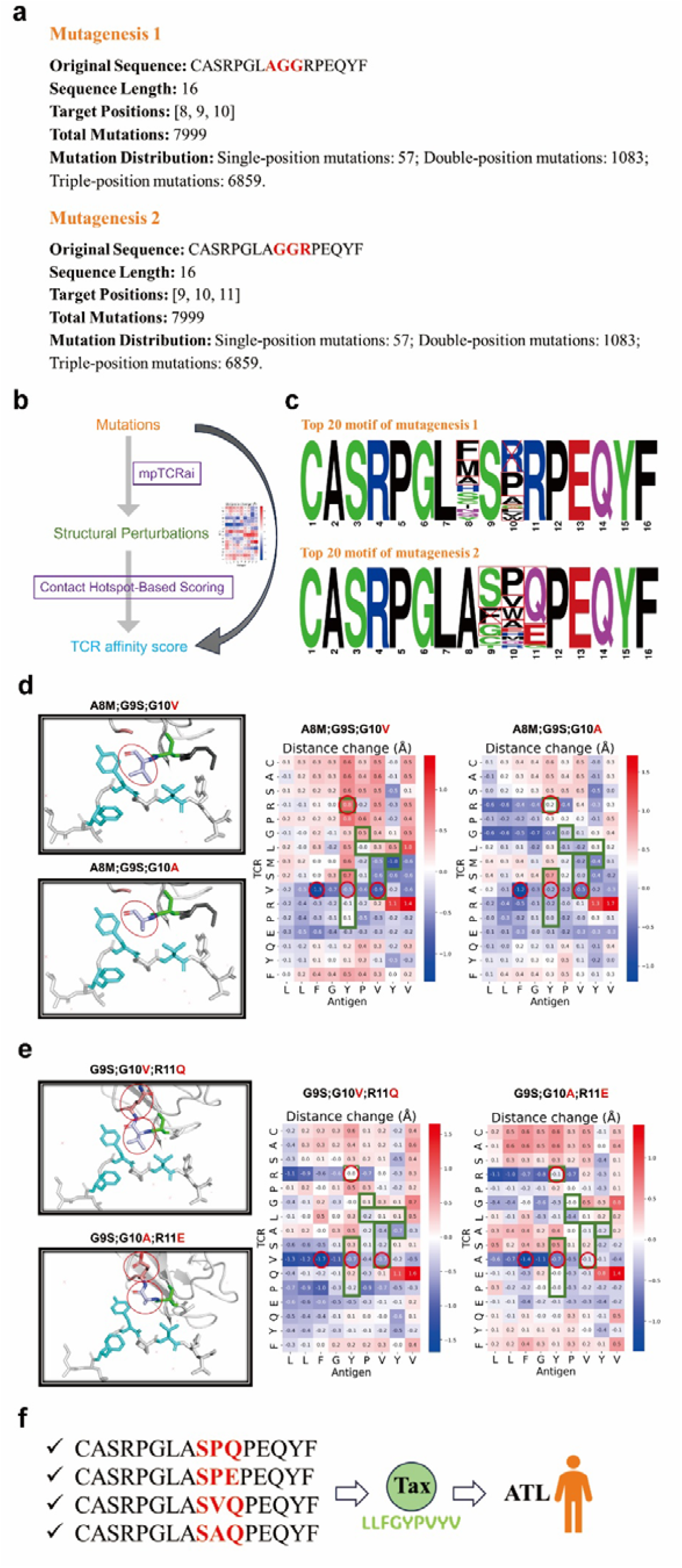
Rational engineering of high-affinity A6-TCR variants for ATL therapy via large-scale in silico screening. **a**. Combinatorial library design strategy. Two comprehensive mutant libraries were constructed by targeting critical CDR3 residues: Library 1 (positions 8-10, 7,999 variants) and Library 2 (positions 9-11, 7,999 variants). **b**. High-throughput in silico prioritization. The mpTCRai framework rapidly profiled the structural and energetic landscapes of all 15,998 variants, enabling the precise filtering of top candidates based on predicted affinity. **c**. Emergent sequence motifs. Sequence logos reveal distinct, position-specific amino acid preferences for Library 1 (upper) and Library 2 (lower) among high-ranking variants. **d**. Structural impact of 8–9–10 region mutations. Structural analysis reveals a deleterious trade-off: while the A-to-V substitution reduces the distance to the hydrophobic pocket, it disrupts the critical hydrogen-bond network with Arginine-4, leading to destabilization. **e**. Structural advantage of 9–10–11 region mutations. A representative top candidate demonstrates dual-optimization: simultaneous interfacial compaction (reduced distance) and preservation of the key hydrogen-bond anchor with residue 4, confirming a robust binding mechanism. **f**. Identification of therapeutic leads. Systematic screening yielded four novel TCR variants with predicted affinities superior to the original positive control, establishing them as promising preclinical candidates for ATL immunotherapy.

We then applied the mpTCRai model to analyze the combined set of 15,998 mutant TCRs (**Fig. 6b**). By predicting structural perturbations and evaluating binding affinity potential to the Tax epitope, we assigned affinity scores to each mutant and selected top 20 candidates from each library (**Table 2 and Table 3**). The final entry in blue represents a positive control with known enhanced TCR activation capacity.

**Table 2.** Top 20 mutants of A6-TCR in positions 8, 9, 10.

| Rank | Mutant ID | Mutated Sequence | Mutation Sites | Positions | Original AAs | Mutated AAs | Score |
| --- | --- | --- | --- | --- | --- | --- | --- |
| 1 | mutant_006462 | CASRPGLSSRRPEQYF | A8S;G9S;G10R | 8;9;10 | A;G;G | S;S;R | 11.36433 |
| 2 | mutant_006477 | CASRPGLSSWRPEQYF | A8S;G9S;G10W | 8;9;10 | A;G;G | S;S;W | 10.93735 |
| 3 | mutant_00<br>5752 | CASRPGLFS<br>PRPEQYF | A8F;G9S;G10P | 8;9;10 | A;G;G | F;S;P | 10.87494 |
| 4 | mutant_00<br>5740 | CASRPGLFS<br>RRPEQYF | A8F;G9S;G10R | 8;9;10 | A;G;G | F;S;R | 9.598985 |
| 5 | mutant_00<br>3935 | CASRPGLHS<br>RRPEQYF | A8H;G9S;G10R | 8;9;10 | A;G;G | H;S;R | 9.531574 |
| 6 | mutant_00<br>5391 | CASRPGLMS<br>PRPEQYF | A8M;G9S;G10P | 8;9;10 | A;G;G | M;S;P | 9.33271 |
| 7 | mutant_00<br>3947 | CASRPGLHS<br>PRPEQYF | A8H;G9S;G10P | 8;9;10 | A;G;G | H;S;P | 9.198351 |
| 8 | mutant_00<br>5748 | CASRPGLFS<br>LRPEQYF | A8F;G9S;G10L | 8;9;10 | A;G;G | F;S;L | 9.025557 |
| 9 | mutant_00<br>5743 | CASRPGLFS<br>CRPEQYF | A8F;G9S;G10C | 8;9;10 | A;G;G | F;S;C | 8.691485 |
| 10 | mutant_00<br>5396 | CASRPGLMS<br>VRPEQYF | A8M;G9S;G10V | 8;9;10 | A;G;G | M;S;V | 8.66472 |
| 11 | mutant_00<br>1047 | CASRPGLAS<br>RRPEQYF | G9S;G10R | 9;10 | G;G | S;R | 8.655459 |
| 12 | mutant_00<br>5379 | CASRPGLMS<br>RRPEQYF | A8M;G9S;G10R | 8;9;10 | A;G;G | M;S;R | 8.181301 |
| 13 | mutant_00<br>7557 | CASRPGLYS<br>PRPEQYF | A8Y;G9S;G10P | 8;9;10 | A;G;G | Y;S;P | 8.140238 |
| 14 | mutant_00<br>1769 | CASRPGLNS<br>RRPEQYF | A8N;G9S;G10R | 8;9;10 | A;G;G | N;S;R | 7.797006 |
| 15 | mutant_00<br>4308 | CASRPGLISP<br>RPEQYF | A8I;G9S;G10P | 8;9;10 | A;G;G | I;S;P | 7.728935 |
| 16 | mutant_00<br>1059 | CASRPGLAS<br>PRPEQYF | G9S;G10P | 9;10 | G;G | S;P | 7.40728 |
| 17 | mutant_00<br>2491 | CASRPGLCS<br>RRPEQYF | A8C;G9S;G10R | 8;9;10 | A;G;G | C;S;R | 7.389081 |
| 18 | mutant_00<br>2852 | CASRPGLQS<br>RRPEQYF | A8Q;G9S;G10R | 8;9;10 | A;G;G | Q;S;R | 7.252611 |
| 19 | mutant_00<br>5739 | CASRPGLFS<br>ARPEQYF | A8F;G9S;G10A | 8;9;10 | A;G;G | F;S;A | 7.061561 |
| 20 | mutant_00<br>5378* | CASRPGLMS<br>ARPEQYF | A8M;G9S;G10A | 8;9;10 | A;G;G | M;S;A | 7.027033 |

\*The mutant\_005378 is positive control.

**Table 3.**
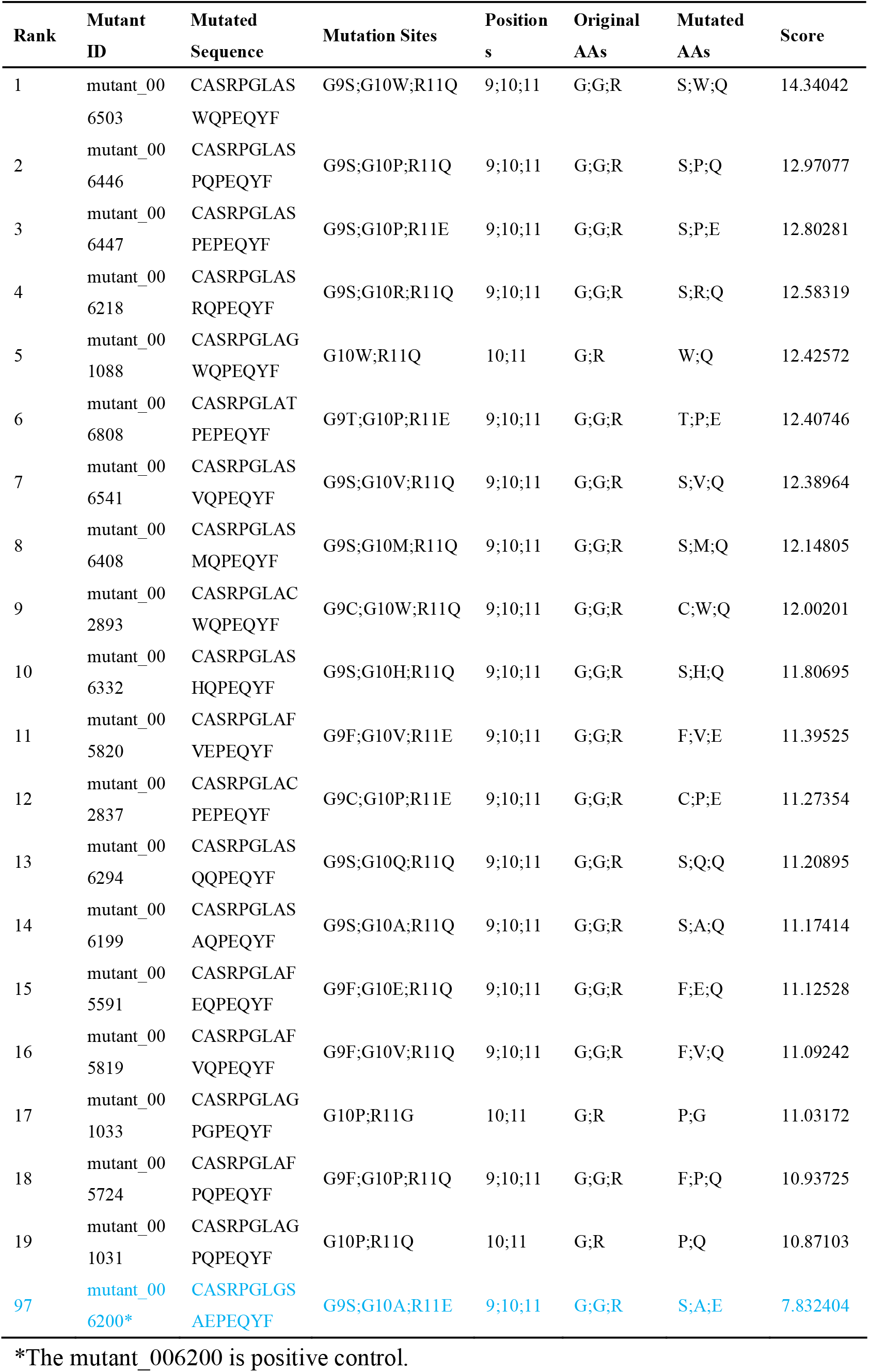
Top 20 mutants of A6-TCR in positions 9, 10, 11.

Subsequent motif analysis revealed distinct residue preferences among the top candidates. For the 8–9–10 mutation set, position 8 favored nonpolar residues such as F or M; position 9 exhibited high specificity, exclusively accommodating the polar residue S; and position 10 showed a strong preference for hydrophobic residues (e.g., P, A, L), consistent with its role in engaging the Tax hydrophobic pocket. For the 9–10–11 set, position 9 again preferred S; position 10 maintained a pronounced bias toward hydrophobic residues (P, V, W, A), reinforcing its central role in hydrophobic interactions; and position 11 tended to be substituted with Q or E (**Fig. 6c**). Based on these patterns, we selected 10 candidate TCR mutants for further validation.

In-depth structural analysis revealed region-specific functional impacts: For the 8–9–10 mutants, substitutions such as A-to-V shortened the distance to the hydrophobic pocket but disrupted the essential hydrogen-bond network involving arginine at position 4 (**Fig. 6d**). This pattern was observed across all other candidates in this group, suggesting that these mutations are likely non-functional or deleterious. Thus, the original positive control sequence (8M-9S-10A) remains optimal for this region. In contrast, for the 9–10–11 mutants, position 10 substitutions reduced the interfacial distance while preserving the key hydrogen bond with residue 4 **(Fig. 6e**), indicating beneficial effects. Through systematic evaluation, we identified four TCR variants predicted to outperform the original positive control (9S-10A-11E). All details of the mutagenesis study on A6-TCR are provided in the **Supplementary file**.

These variants represent promising engineered TCR candidates for the treatment of Tax-associated adult T-cell leukemia **(Fig. 6f**).

## Discussion

To decode specificity of TCR-antigen interaction, we developed mpTCRai, a novel deep learning framework that simulates the two-stage sequential process of T cell activation through dual-representation modeling and MPCA mechanism. A key advancement of mpTCRai lies in its high biological fidelity useful for the integration of structural perturbation with functional and mechanistic insights. Unlike conventional single-step predictors that often overestimate immunogenicity by focusing exclusively on binding affinity^40–43^, our framework recapitulates the two-stage nature of actual T cell activation. Such a design is critical, as recent studies have shown that the majority of MHC-presented antigens cannot to elicit productive immune responses^44–46^. This biological fidelity proves particularly valuable for addressing persistent challenges in TCR engineering and cross-reactivity assessment in computational immunology.

First, the contact hotspot-based scoring function addresses a key gap in TCR–antigen interaction analysis by revealing that functional outcomes depend more on specific residue-level interactions than on global binding energy. This approach achieves a stronger correlation with experimental ΔΔG values (r = –0.88), establishing robust links between sequence features, structural perturbation, and binding affinity. Moreover, it reveals molecular switches that can enhance TCR affinity—such as hydrophobic core formation and conformational stabilization—deepening the knowledge about immune recognition for actionable guidance in rational TCR engineering.

Second, the analysis on Tax Y5 mutations illustrates how the integration of structural and chemical information overcomes the limitations of distance-only metrics. Two key findings support this: (i) the contact hotspot-based scoring function successfully identifies cross-reactive mutants (Y5F and Y5W), confirming its predictive power; (ii) the Y5M mutant exemplifies the risks of relying solely on geometric criteria. The Y5M case is particularly instructive—while distance-based metrics suggest potential cross-reactivity, integrated analysis incorporating chemical properties reveals functional impairment, highlighting the necessity of a multifactorial approach in TCR specificity profiling.

Together, our method has been systematically validated through case studies from both TCR-centric (TCR variants) and pHLA-centric (antigen variants) perspectives. From the TCR-centric perspective, our approach accurately identifies a key molecular switch that regulate TCR activation (**Fig. 4**). From the antigen-centric perspective, it effectively detects peptide variants exhibiting cross-reactive potential (**Fig. 5**). Based on these findings, we performed computational multi-site combinatorial mutagenesis on the A6-TCR, proposing four mutant variants to guide subsequent in vitro/vivo validation for ATL (**Fig. 6**). This integrated framework thus establishes a robust technical foundation for constructing an efficient pipeline encompassing mutation design, TCR affinity assessment, cross-reactivity evaluation, and clinical candidate selection, with strong potential to accelerate the preclinical development of neoantigen vaccines and TCR-T cell therapies (**Fig. S3b**).

Despite these advances, several limitations of mpTCRai must be acknowledged. First, like all data-driven methods, the model’s performance is inherently limited by the availability and representativeness of structural data. As shown in **Fig. S4b and c**, the model exhibits reduced accuracy for sequences with atypical lengths or rare structural motifs that are underrepresented in the training set. Furthermore, the results in **Fig. S4d** indicate that predictive performance is primarily constrained by biases in the training data distribution rather than by fundamental algorithmic limitations. This finding underscores the value of our newly curated dataset—currently the largest available structural resource for HLA–antigen–TCR ternary complexes—which will be made publicly available to facilitate broader performance evaluations in computational immunology.

Second, it is important to note the limitations regarding the generalizability of our ΔΔG analysis. Currently, the proposed mapping from structural perturbations to energetic changes is validated solely within the A6-TCR system due to the limited availability of comprehensive experimental thermodynamic data for other complexes. Therefore, this analysis serves primarily as a proof-of-concept demonstrating the potential downstream utility of mpTCRai’s structural predictions. Extending this approach to formulate a universal predictive rule will necessitate rigorous evaluation across diverse TCR-antigen pairs as more experimental thermodynamic measurements become available.

Finally, the current framework treats residues as static entities. To ensure robust cross-reactivity and affinity predictions across broader immunological contexts, future iterations must transcend purely geometric predictions. Incorporating complementary physics-based approaches, such as molecular dynamics (MD) simulations and physics-based energy functions, could better capture the conformational flexibility and time-resolved intermolecular interactions essential to accurate TCR recognition modeling.

## Methods

### Data processing

Two HLA-antigen-TCR ternary structure datasets are used by this work: the STCRDab dataset^24^ containing 122 complexes and Feng’s dataset^47^ comprising 86 complexes. Initial quality assessment found data redundancy between these two data resources. Specifically, utilizing the STCRDab database as the baseline, we applied a sequential filtering process. First, we retained only structural data representing complete TCR-peptide-MHC ternary complexes. Next, we restricted the samples to those containing human-derived MHC molecules. Following this, we computed the residue distance matrix between the TCR and the antigen. Samples were excluded if the distance matrix could not be computed or if all pairwise residue distances exceeded 5 Å. This standardized filtering pipeline yielded the final training dataset of 90 samples (the training dataset, **Supplementary Table 1**). By applying identical data curation standards to Feng’s dataset and performing strict de-duplication against our training set, we retained only 7 test samples (the test1 dataset, **Supplementary Table 2**).

As these two datasets were published prior to 2017, we conducted an extensive search at Protein Data Bank (PDB)^48^ to identify and incorporate post-2017 ternary structures, thereby building a novel third dataset (n=67, named test2 dataset, **Supplementary Table 3**). Briefly, we systematically queried PDB in May 2025 using the following strict conditions: (1) Text search inside all attributes = “TCR” AND “antigen”; (2) Scientific Name of the Source Organism = “Homo sapiens”; (3) Release Date = [2017-09-01 to 2029-12-31].

Each input entry for mpTCRai consist of three key components: full-length HLA sequences (from the IPD-IMGT/HLA database^49^), antigen peptides, and TCR CDR3β sequences (from 10 to 20 amino acids).

We downloaded the three-dimensional structural coordinates of all samples from the PDB database using Biopython^50^ (version: 1.85). For each HLA-antigen-TCR complex, we calculated the minimum inter-residue distances between the TCR and antigen sequences to generate a TCR-antigen residue distance matrix. Following the same criteria as TEIM^18^, we defined residue pairs with distances ≤5□□ as contact residues and constructed its corresponding binary contact matrix. This contact matrix also serves as the ground truth for model training and evaluation, where a value of 1 indicates a contact residue pair and 0 represents non-contact residues.

### Model architecture

For vector representations of the sequences, we employed the ESM2^19^ (650M) protein language model to encode HLA and antigen sequences, while utilizing TCR-BERT^20^ for TCR sequence embedding.

To simulate the presentation process of antigens, we implemented three distinct attention learning methods: self-attention^51^, co-attention^52^, and cross-attention^51,53^ (with antigens as query) to capture the interactions between HLAs and antigens, ultimately generating the pHLA complex representation.

For the modeling of T cell recognition, we similarly applied self-attention, co-attention, and a novel mutual-perspective cross-attention mechanisms to simulate TCR-pHLA interactions.

Let 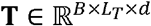 and 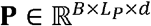 be the input representation for the TCR and pHLA, respectively. The model computes a mutual-perspective cross-attention ℱ_interaction_ through the following sequential operations:

#### Step 1: TCR-Centric Attention and Projection

The TCR representation is enriched using the pHLA context as the source of keys and values.

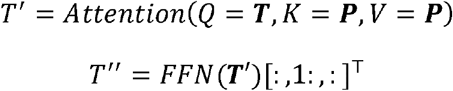

#### Step 2: pHLA-Centric Attention and Projection

Symmetrically, the pHLA representation is enriched using the TCR context.

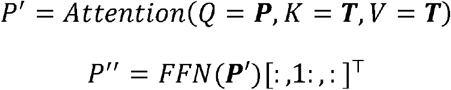

#### Step 3: Construction of the Paired-Feature Tensors

The refined representations are broadcast to create 4D tensors encompassing all residue pairs.

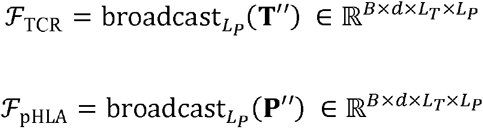

#### Step 4: Generation of the mutual perspective map

The final interaction map is formed by fusing the two feature tensors via element-wise multiplication.

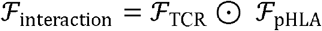

The additive loss function was formulated for the two prediction tasks. For the regression task (prediction of the residue distance matrix), we employed Mean Squared Error (MSE) Loss, while for the classification task (prediction of the residue contact matrix), we used Binary Cross-Entropy (BCE) Loss. These two loss functions were jointly optimized during training, given as

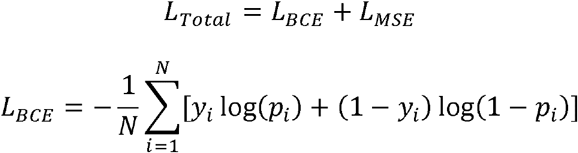

where, *y*_*i*_ stands for ground truth contact label (1 if residues ≤5 Å, else 0); *p*_*i*_ means a predicted contact probability.

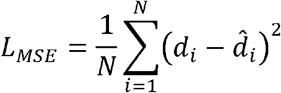

where, *d*_*i*_ represents true distance (from PDB); 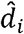 represents a predicted distance.

To efficiently fine-tune the ESM2 and TCR-BERT models, we applied LoRA technology. Training was done using an AdamW optimizer^54^ with a learning rate of 5e-5, on an Nvidia RTX4090 GPU.

### Performance evaluation metrics

To evaluate the predicted TCR-antigen distance results by mpTCRai, we calculated Pearson correlation coefficients between these predicted and the actual residue-pair distances.

Following the same criteria as TEIM, residue pairs within 5□□ were defined as contact pairs^18^. We constructed confusion matrices to analyze the distribution of the predicted versus the actual contacting residues and computed four performance metrics: Accuracy, Precision, Recall, and F1-score.

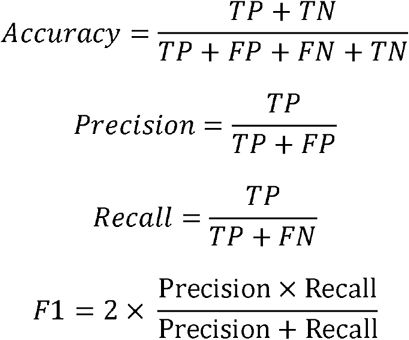

TP: True Positive; FP: False Positive; FN: False Negative; TN: True Negative.

Particularly, Recall specifically measures the model’s ability to capture the minority class (contact residues), which is important for assessing prediction sensitivity at interaction interfaces.

### Hotspot-based scoring function

We proposed a hotspot-based scoring function for enhancing the stratification of functional variants.

Let *M* ∈ *R*^*m*×*n*^ be the original distance matrix, *M*^′^ ∈ *R*^*m*×*n*^ be the comparison matrix; *Δ* = *M*^′^ − *M* be the difference matrix; ℋ = {(*i, j*) | *M*_*ij*_ ≤ 5} denote contact residues; C = {(*i, j*) | 5 < *M*_*ij*_ ≤ 7} represent candidate residues; *Δ*_*ij*_ be the distance change at position (i,j). We define a hotspot-based score S as

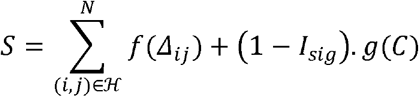

satisfying

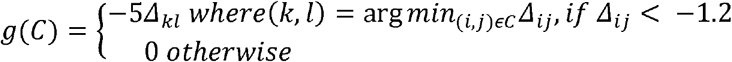

where,

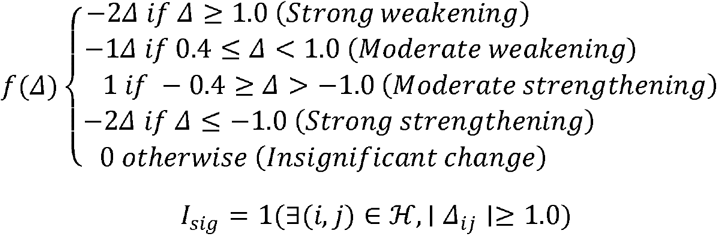

## Acknowledgements

We would like to thank Servier Medical Art for providing the materials used to create the figures.

## Author contributions

Conceptualization: SFL, NYS, JYL; Methodology: SFL, SMZ, SY; Investigation: SFL, JJL, JXC; Visualization: SFL, SMZ; Supervision: NYS, YP, JYL; Writing-original draft: SFL; Writing-review & editing: NYS, JYL.

## Data availability

The processed data for our model training and evaluation are all available at GitHub repository website https://anonymous.4open.science/r/mpTCRai-1ED1AI. The raw data were all downloaded from public websites: PDB database (https://www.rcsb.org/); IPD-IMGT/HLA database (https://www.ebi.ac.uk/ipd/imgt/hla/).

