## Supplemental materials for "Deep mapping of structural perturbations to energetics enables precise TCR design"

### Supplementary materials

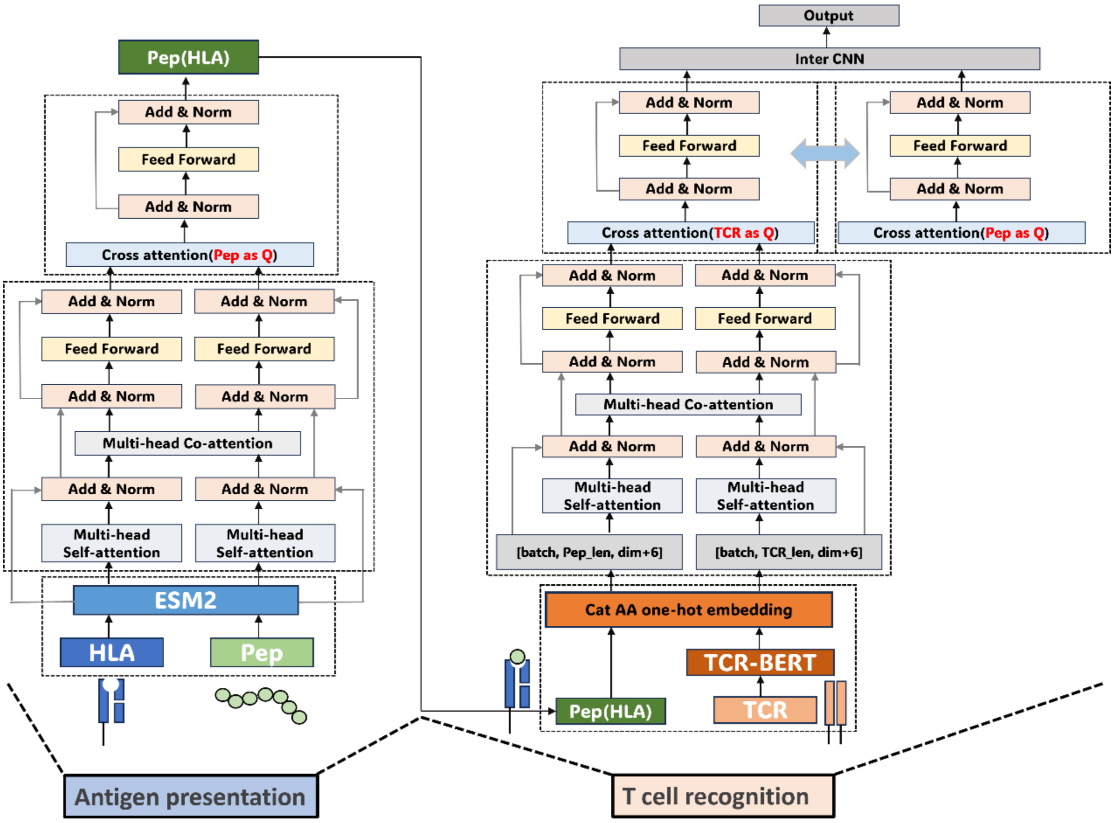


**Figure S1. A detailed schematic data flow of our mutual perspective modeling approach.** The mpTCRai model combines ESM2-based representations for HLA molecules and antigens with TCR-BERT-derived embeddings for TCR sequences. These representations are processed through multi-type attention mechanisms to simulate the two key biological processes: antigen presentation and T cell recognition.

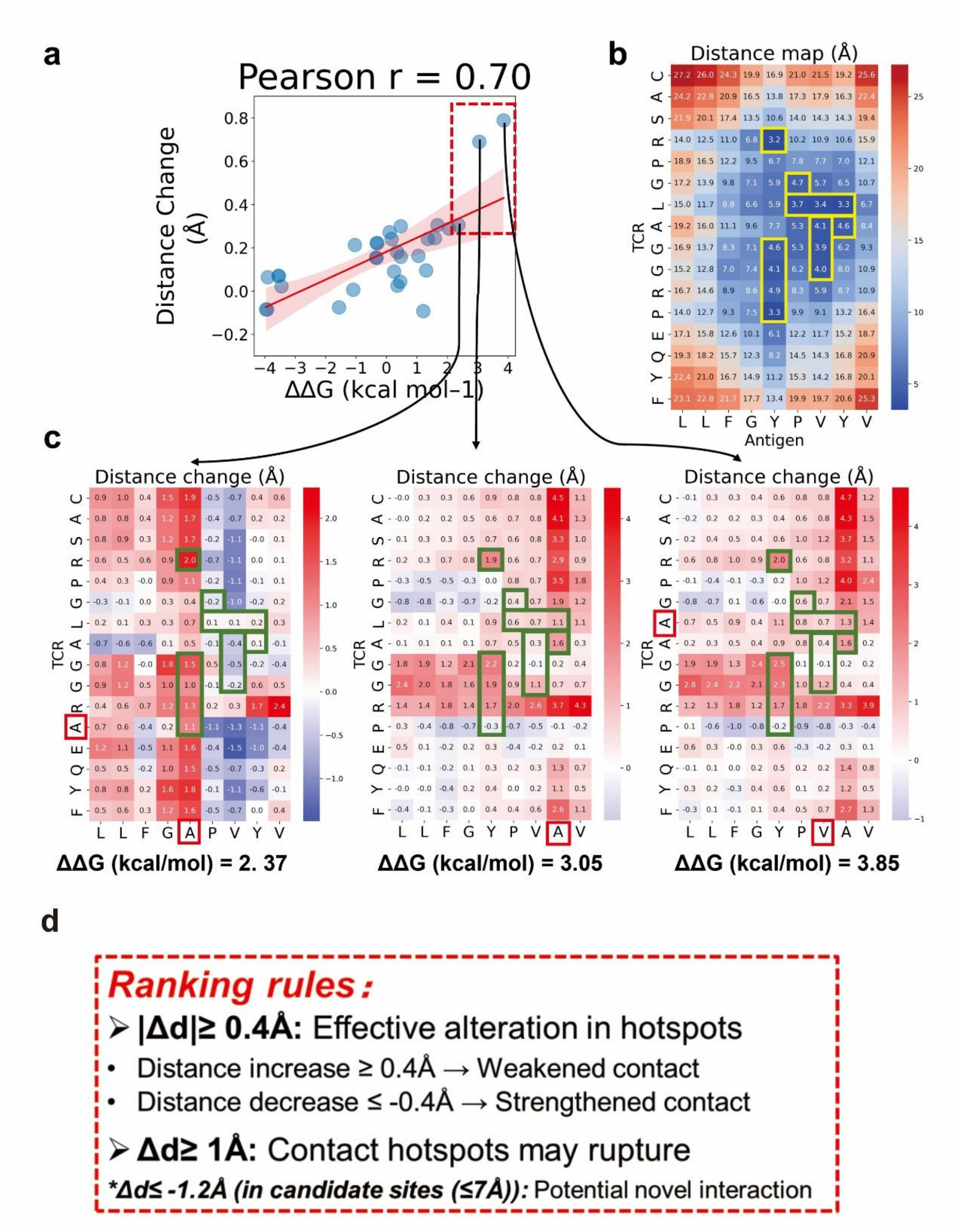


**Figure S2. Structural and energetic analysis of mutation effects using mpTCRai.** **a.** Correlation between predicted structural perturbations (residue distance changes) and experimental binding affinity changes (ΔΔG). **b.** Contact hotspots (yellow box, defined as residue pairs within ≤5 Å) illustrated on the A6-Tax distance map. **c.** Heatmap of residue distance changes for mutations exhibiting significant affinity loss, highlighting disrupted interaction patterns. **d.** Schematic of the scoring rules for the contact hotspot-based function. Heatmap interpretation in *c*: Blue indicates distance reduction (potential interaction strengthening); red indicates distance increase (potential interaction weakening).

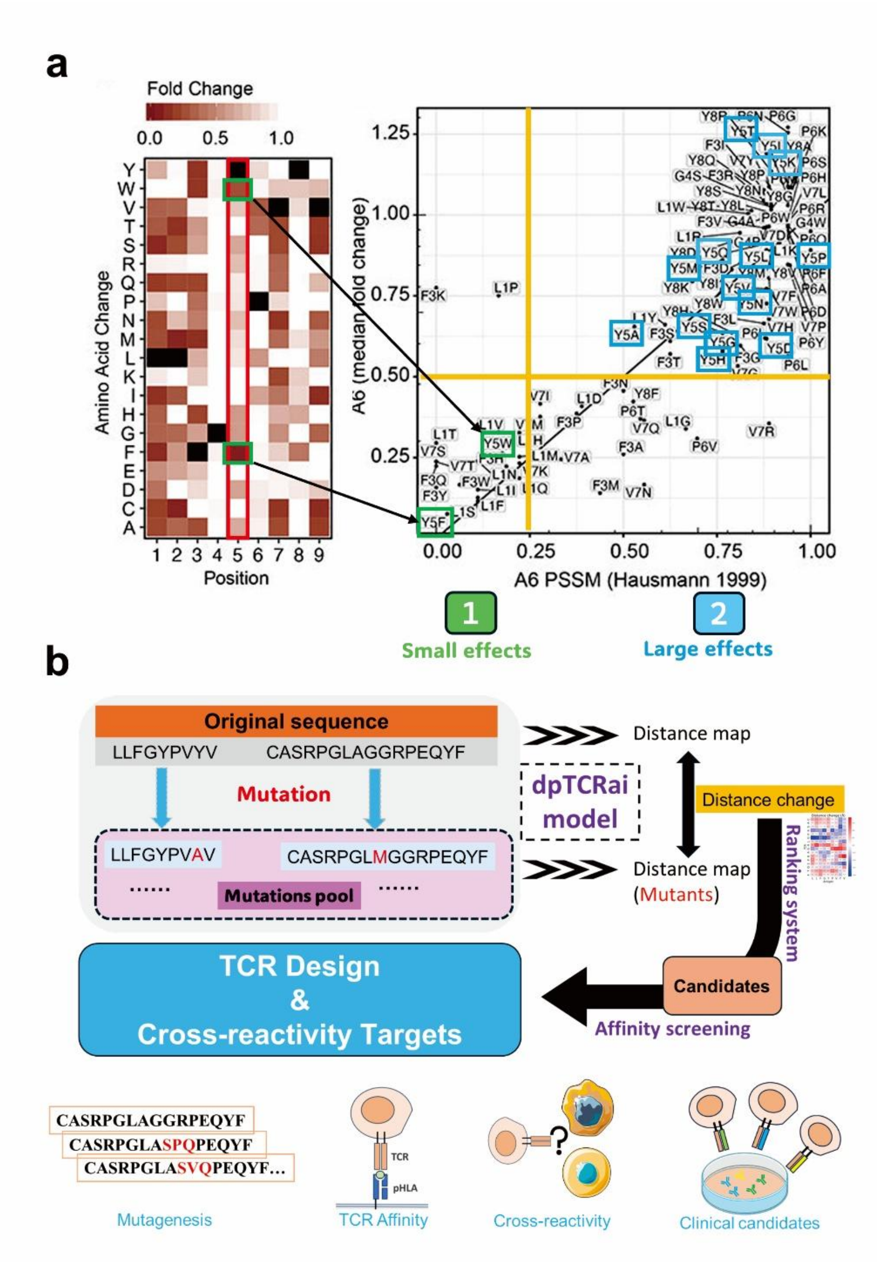


**Figure S3. Classification of cross-reactive substitutions and model applications.** **a.** Activity measurements of Tax Y5 substitutions comparing aromatic-preserving "small effect" variants (Y5F/W) with disruptive "large effect"[37]. **b.** Schematic workflow of the clinical translation pipeline integrating mpTCRai predictions for TCR engineering and neoantigen screening prediction.

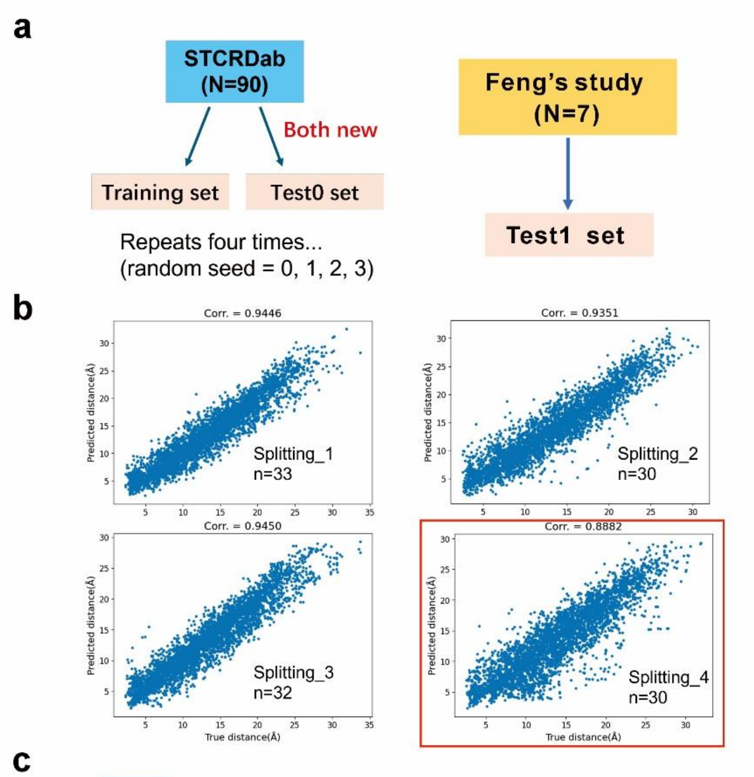


c

| Coef (target_1) | MAE(target_1) | MAPE(target_1) | Roc_auc_score(target_2) | CDR3β | Antigen |
|---|---|---|---|---|---|
| 0.9721 | 0.8147 | 0.0654 | 0.9581 | CASSYVGNTGELFF | SLLMWITQV |
| 0.9670 | 0.9738 | 0.0797 | 0.9494 | CASSYVGNTGELFF | SLLMWITQC |
| 0.9667 | 1.0023 | 0.0748 | 0.9518 | CASSYLGNTGELFF | SLLMWITQC |
| 0.7895 | 2.8940 | 0.1763 | 0.8000 | CASSANPGDSSNEKLFF | VMAPRTLIL |
| 0.8005 | 2.7377 | 0.1754 | 0.7852 | CASSANPGDSSNEKLFF | VMAPRTLVL |
| 0.9383 | 2.3189 | 0.1398 | 0.9281 | CASSQDRDTQYF | VMAPRTLIL |
| 0.9681 | 1.3351 | 0.1060 | 0.9800 | CASSLSFGTEAFF | ELAGIGILTV |
| 0.9653 | 1.0870 | 0.0884 | 0.9701 | CASSLSFGTEAFF | AAGIGILTV |
| 0.9607 | 1.4347 | 0.1078 | 0.9572 | CASSLSFGTEAFF | SMLGIGIVPV |
| 0.8601 | 2.8634 | 0.1941 | 0.9103 | CASSLSFGTEAFF | MMWDRGLGMM |
| 0.9762 | 1.0695 | 0.0846 | 0.9675 | CASSWSFGTEAFF | ELAGIGILTV |
| 0.9227 | 1.9744 | 0.1600 | 0.9408 | CASSWSFGTEAFF | AAGIGILTV |
| 0.9893 | 0.5729 | 0.0447 | 1.0000 | CAISEVGVGQPQHF | AAGIGILTV |
| 0.9813 | 0.5203 | 0.0565 | 0.9995 | CAWSETGLGTGELFF | AAGIGILTV |
| 0.9786 | 0.5388 | 0.0584 | 0.9970 | CAWSETGLGMGGWQF | AAGIGILTV |
| 0.8694 | 2.6384 | 0.1744 | 0.9417 | CASSARSGELFF | HPVGEADYFEY |
| 0.8403 | 2.5921 | 0.1862 | 0.9150 | CASSARSGELFF | HPVGDADYFEY |
| 0.8756 | 2.6630 | 0.1713 | 0.9386 | CASSARSGELFF | HPVGQADYFEY |
| 0.9276 | 1.6293 | 0.1047 | 0.9359 | CASSLWEKLAKNIQYF | ALWGPDPAAA |
| 0.9305 | 1.4550 | 0.1057 | 0.9439 | CASSLWEKLAKNIQYF | YQFGPDFPIA |
| 0.9307 | 1.4562 | 0.1059 | 0.9247 | CASSLWEKLAKNIQYF | RQWGPDPAAV |
| 0.9252 | 1.5979 | 0.1176 | 0.9958 | CASSLWEKLAKNIQYF | YLGGPDFPTI |
| 0.9307 | 1.6792 | 0.1164 | 0.9277 | CASSLWEKLAKNIQYF | MVWGPDPLYV |
| 0.9220 | 1.4940 | 0.1068 | 0.9343 | CASSLWEKLAKNIQYF | RQFGPDFPTI |
| 0.9319 | 1.2292 | 0.0954 | 0.9247 | CASSLWEKLAKNIQYF | AQWGPDPAAA |
| 0.9007 | 2.1561 | 0.1577 | 0.9379 | CASTGSYGYTF | KAFSPEVIPMF |
| 0.9282 | 1.8339 | 0.1415 | 0.9163 | CASRYRDDSYNEQFF | EENLLDFVRF |
| 0.5977 | 3.7104 | 0.2798 | 0.6753 | CASSFTWTSGGATDTQYF | FLRGRAYGL |
| 0.9269 | 1.9877 | 0.1530 | 0.9640 | CASSLGQAYEQYF | FLRGRAYGL |
| 0.9302 | 1.6370 | 0.1306 | 0.9667 | CASGQGNFDIQYF | FLRGRAYGL |

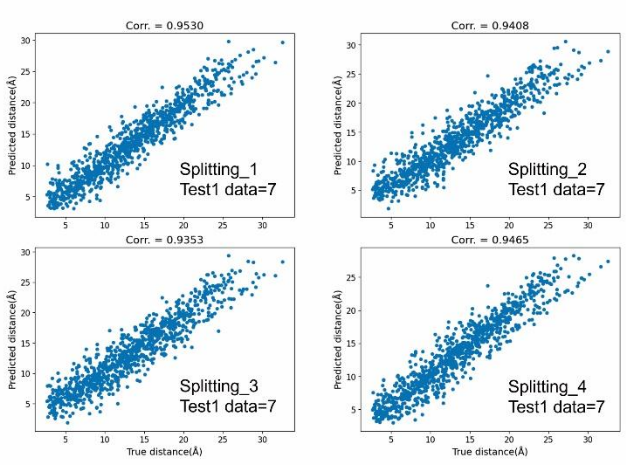


**Figure S4. Evaluation of data dependency and prediction consistency across random partitions. a.** Data partitioning strategy. Schematic illustrating the randomized splitting approach used to generate four independent training/validation partitions. **b.** Model performance consistency across partitions. Dot plot showing Pearson correlation coefficients for residue distance prediction across four distinct data partitions. **c.** Sample-level prediction accuracy in Partition 4. Distribution of Pearson correlation values per test sample (blue boxes), with outliers (coefficient < 0.9) highlighted in yellow. **d.** Validation on an external dataset. Performance comparison of four model instances (trained with different random seeds) on Feng's validation set (n = 7), demonstrating high consistency (r > 0.93) across all replicates**.**

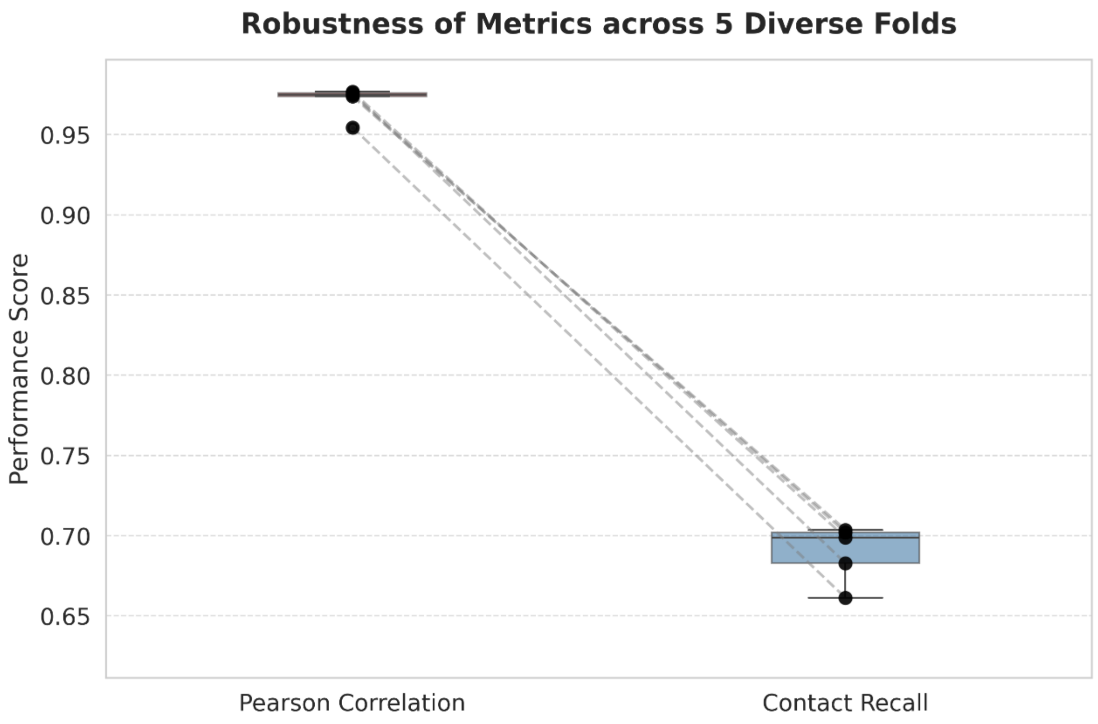


**Figure S5. Figure S5. Robustness evaluation of mpTCRai through 5-fold cross-validation on the expanded dataset.** Boxplots illustrate the predictive performance of the model across five independent folds, using a combined dataset of 164 structurally characterized HLA-antigen-TCR ternary complexes (N≈131 per training fold). Two key metrics are evaluated: the Pearson correlation of TCR-antigen residue distances and the contact residue recall. The results demonstrate highly stable predictive capability, with the residue distance correlation consistently ranging between 0.95 and 0.98, and the contact recall ranging from 0.66 to 0.70 across all folds.

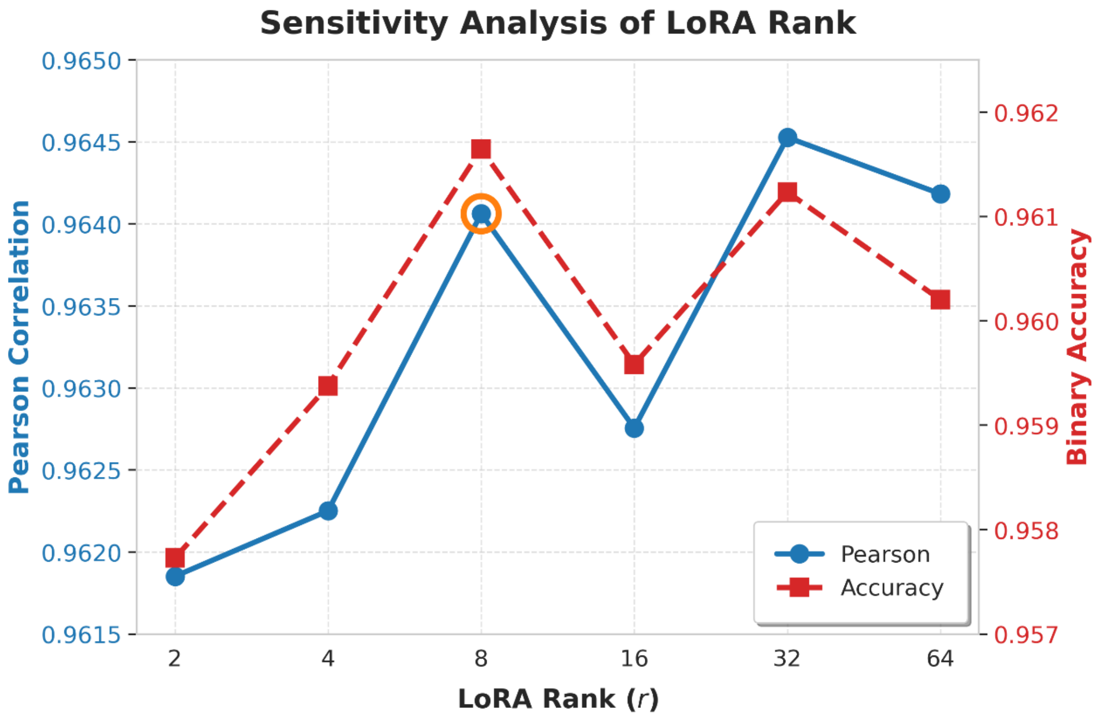


**Figure S6. Sensitivity analysis of the LoRA rank hyperparameter.** A dual-axis line chart illustrating the model's predictive performance across various LoRA rank configurations, evaluated using the 90-sample training dataset (sampled from the combined dataset of 164). The left y-axis denotes the Pearson correlation coefficient for TCR-antigen residue distance prediction (ranging from 0.9615 to 0.9650), while the right y-axis indicates the contact prediction accuracy (ranging from 0.957 to 0.962). The extremely narrow variation in both metrics demonstrates that the specific LoRA rank does not significantly impact overall model robustness. However, an inflection point is observed at a rank of 8, indicating an optimal trade-off between model complexity and predictive efficiency, thus justifying its selection as the default hyperparameter in our framework.

### Analysis Report

Mutagenesis study on A6-TCR

#### 1. Introduction

This report summarizes the results of a computational mutagenesis study on the TCR sequence CASRPGLAGGRPEQYF. The optimization strategy is guided by prior structural analyses, which revealed that the recognition of the Tax epitope by the optimized A6-TCR relies on two critical structural-functional relationships: first, the interaction between residue **100A** on the CDR3 loop and the hydrophobic pocket of the Tax epitope; and second, a key hydrogen bond between residue **94R** on the CDR3 loop and the **Y5** residue of the Tax epitope. Building on this foundation, our deep learning model was used to systematically evaluate comprehensive mutations at positions 8, 9, and 10 (positions 9, 10, and 11)—located within the CDR3 loop—to identify variants that potentially enhance these critical interactions and improve overall binding affinity. The analysis includes mutation profiling, affinity scoring, motif identification, and candidate sequence evaluation.

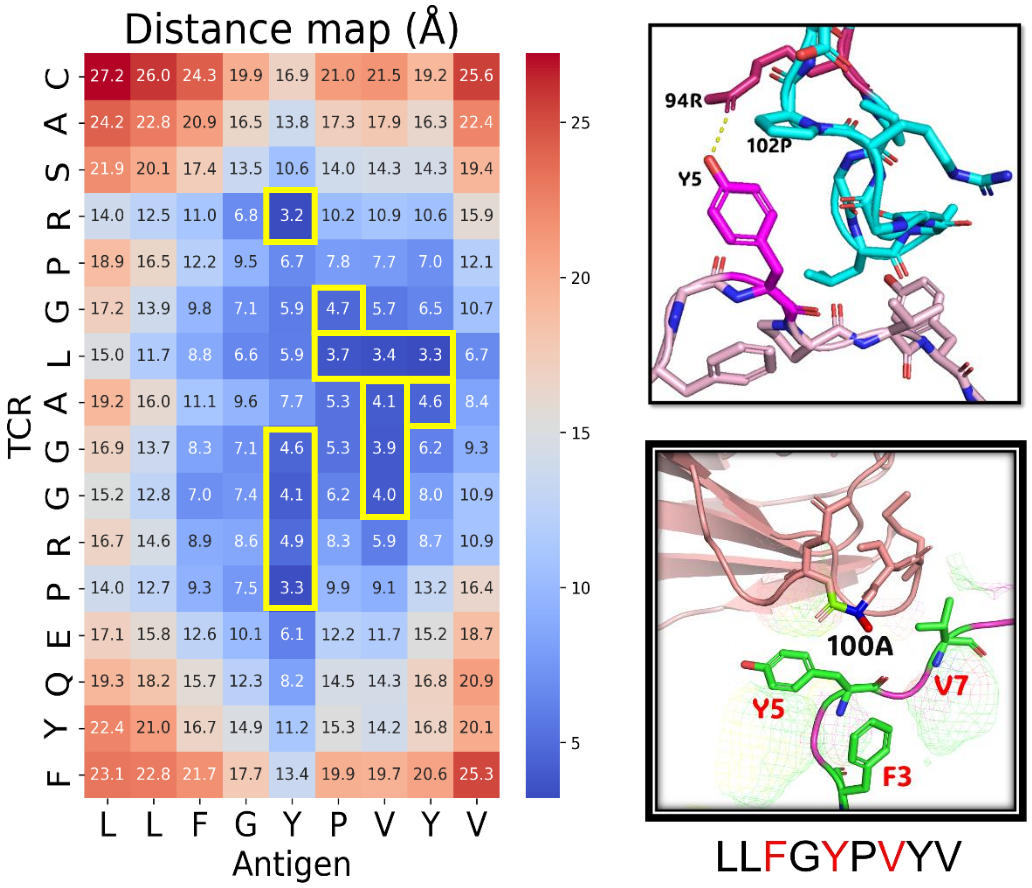


#### 2. Mutation Overview

##### 2.1. Target Positions: 8, 9, 10

**Original Sequence:** CASRPGL**AGG**RPEQYF

**Sequence Length:** 16

**Target Positions:** [8, 9, 10]

**Total Mutations:** 7999

**Mutation Distribution:** Single-position mutations: 57; Double-position mutations: 1083; Triple-position mutations: 6859.

##### 2.2. Target Positions: 9, 10, 11

**Original Sequence:** CASRPGLAGGRPEQYF

**Sequence Length:** 16

**Target Positions:** [9, 10, 11]

**Total Mutations:** 7999

**Mutation Distribution:** Single-position mutations: 57; Double-position mutations: 1083; Triple-position mutations: 6859.

#### 3. Top 20 Mutants by Hotspot-Based Scoring Approach

##### 3.1 Target Positions: 8, 9, 10

- **Ranking list**

| Rank | Mutant_ID | Mutated_Sequence | Mutation_Sites | Positions | Original_AAs | Mutated_AAs | Score |
|---|---|---|---|---|---|---|---|
| 1 | mutant_006462 | CASRPGLSSRRPEQYF | A8S;G9S;G10R | 8;9;10 | A;G;G | S;S;R | 11.36433 |
| 2 | mutant_006477 | CASRPGLSSWRPEQYF | A8S;G9S;G10W | 8;9;10 | A;G;G | S;S;W | 10.93735 |
| 3 | mutant_005752 | CASRPGLFSPRPEQYF | A8F;G9S;G10P | 8;9;10 | A;G;G | F;S;P | 10.87494 |
| 4 | mutant_005740 | CASRPGLFSRRPEQYF | A8F;G9S;G10R | 8;9;10 | A;G;G | F;S;R | 9.598985 |
| 5 | mutant_003935 | CASRPGLHSRRPEQYF | A8H;G9S;G10R | 8;9;10 | A;G;G | H;S;R | 9.531574 |
| 6 | mutant_005391 | CASRPGLMSPRPEQYF | A8M;G9S;G10P | 8;9;10 | A;G;G | M;S;P | 9.33271 |
| 7 | mutant_003947 | CASRPGLHSPRPEQYF | A8H;G9S;G10P | 8;9;10 | A;G;G | H;S;P | 9.198351 |
| 8 | mutant_005748 | CASRPGLFSLRPEQYF | A8F;G9S;G10L | 8;9;10 | A;G;G | F;S;L | 9.025557 |
| 9 | mutant_005743 | CASRPGLFSCRPEQYF | A8F;G9S;G10C | 8;9;10 | A;G;G | F;S;C | 8.691485 |
| 10 | mutant_005396 | CASRPGLMSVRPEQYF | A8M;G9S;G10V | 8;9;10 | A;G;G | M;S;V | 8.66472 |
| 11 | mutant_001047 | CASRPGLASRRPEQYF | G9S;G10R | 9;10 | G;G | S;R | 8.655459 |
| 12 | mutant_005379 | CASRPGLMSRRPEQYF | A8M;G9S;G10R | 8;9;10 | A;G;G | M;S;R | 8.181301 |
| 13 | mutant_007557 | CASRPGLYSPRPEQYF | A8Y;G9S;G10P | 8;9;10 | A;G;G | Y;S;P | 8.140238 |
| 14 | mutant_001769 | CASRPGLNSRRPEQYF | A8N;G9S;G10R | 8;9;10 | A;G;G | N;S;R | 7.797006 |
| 15 | mutant_004308 | CASRPGLISPRPEQYF | A8I;G9S;G10P | 8;9;10 | A;G;G | I;S;P | 7.728935 |
| 16 | mutant_001059 | CASRPGLASPRPEQYF | G9S;G10P | 9;10 | G;G | S;P | 7.40728 |
| 17 | mutant_002491 | CASRPGLCSRRPEQYF | A8C;G9S;G10R | 8;9;10 | A;G;G | C;S;R | 7.389081 |
| 18 | mutant_002852 | CASRPGLQSRRPEQYF | A8Q;G9S;G10R | 8;9;10 | A;G;G | Q;S;R | 7.252611 |
| 19 | mutant_005739 | CASRPGLFSARPEQYF | A8F;G9S;G10A | 8;9;10 | A;G;G | F;S;A | 7.061561 |
| 20 | mutant_005378* | CASRPGLMSARPEQYF | A8M;G9S;G10A | 8;9;10 | A;G;G | M;S;A | 7.027033 |

***Positive control**: This mutant exhibited a significant enhancement in activity against the Tax peptide, with a calculated ΔΔG of -3.55 kcal/mol.

- **Motif Analysis**

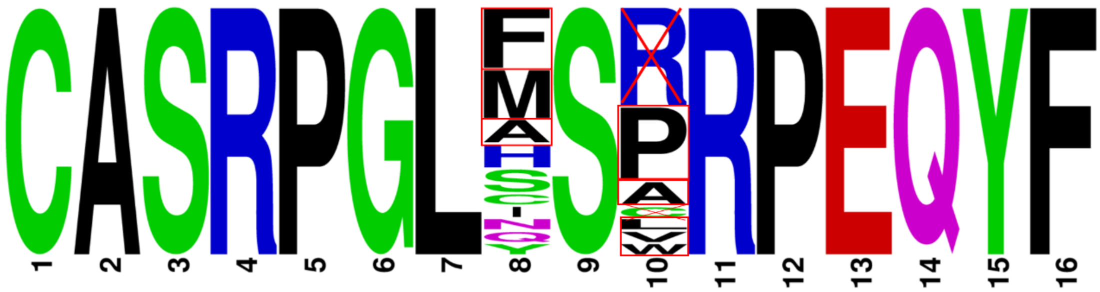


i. Position 8: Strong preference for nonpolar residues (**F, M**)

ii. Position 9: Only preference for polar residues (**S**)

iii. Position 10: Demonstrates a clear inclination towards hydrophobic residues such as **P, A, and L**. This preference is consistent with the residue's role in engaging the Tax hydrophobic pocket. In contrast, while a basic residue like **R** may show a favorable distance score, its charged nature is fundamentally incompatible with the hydrophobic binding environment, making it an unfavorable substitution.

**iv. Conserved Motifs in Top Candidates: [F/M]-[S]-[P/A/L/V]**

#### ◆ Candidate Mutants Evaluation

A8F-G9S-G10P; A8F-G9S-G10A; A8F-G9S-G10L; A8M-G9S-G10P; A8M-G9S-G10V; A8M-G9S-G10A (*positive control).

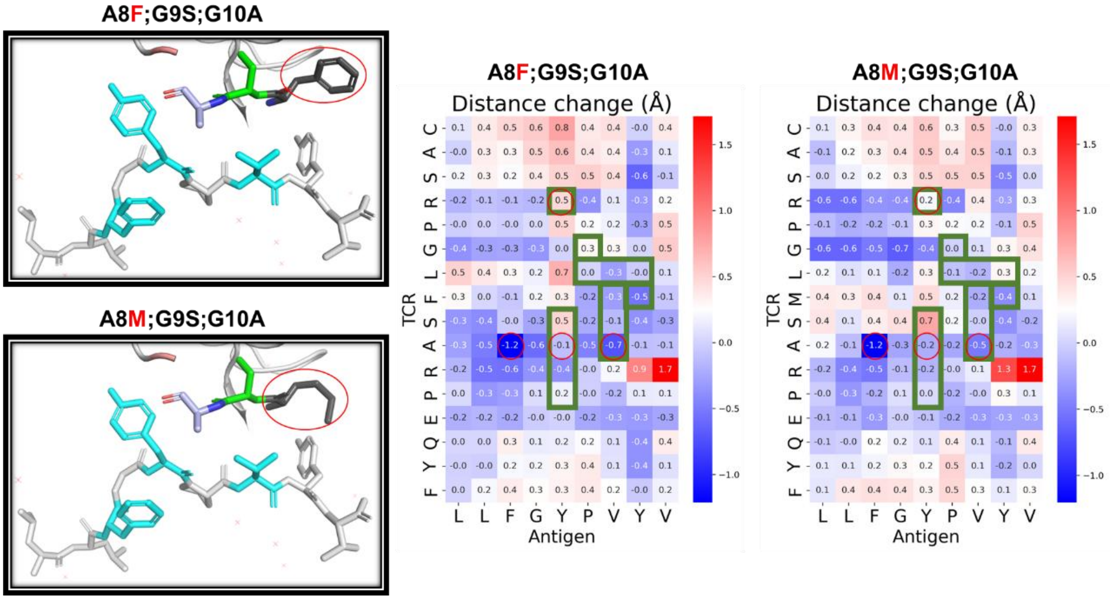


In the MSA positive control, the M8F mutation was found to be functionally detrimental. Despite maintaining the binding state of A10, the mutation compromised a key hydrogen bond with R4, indicating a deleterious effect on structural integrity.

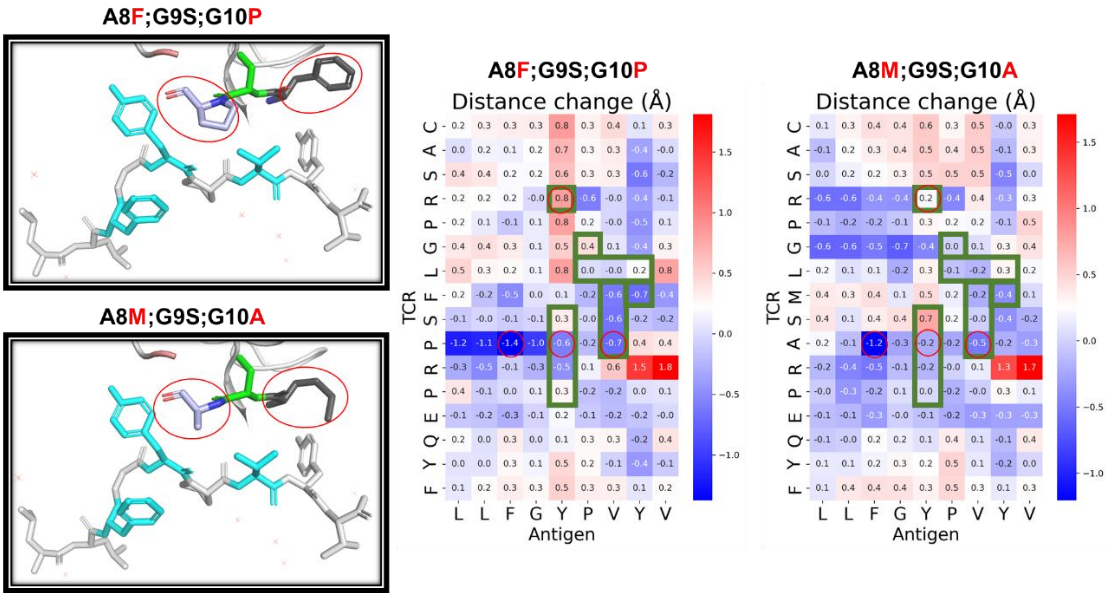


In the MSA positive control, the combined M8F and A10P mutations shortened the binding distance between A10 and the hydrophobic pocket. However, this dual mutation disrupted the critical hydrogen bond mediated by R4. The loss of this key interaction suggests that the mutation is deleterious.

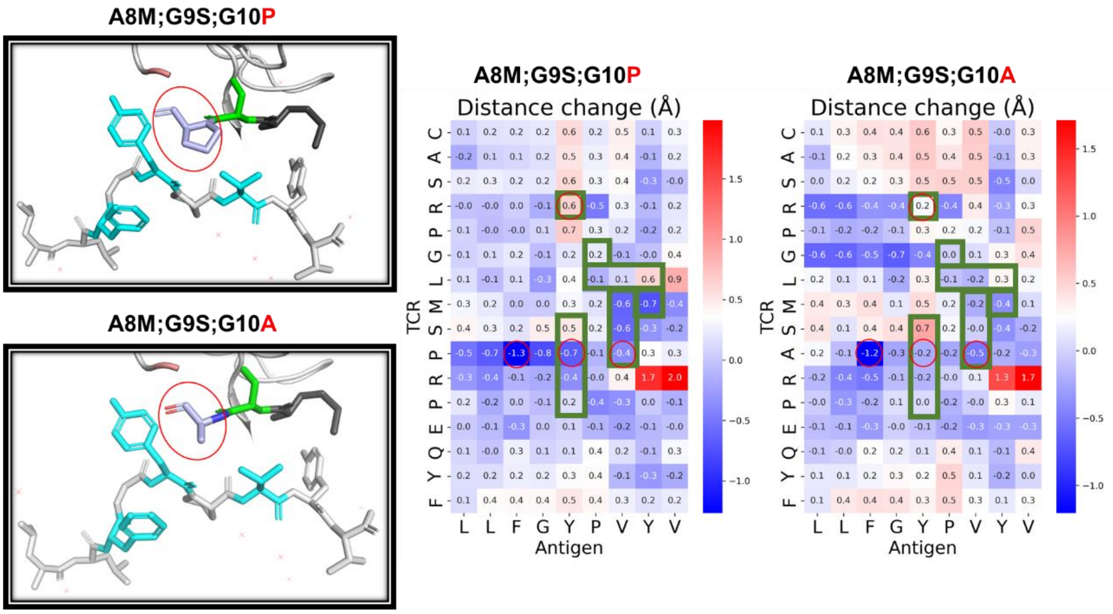


In the MSA positive control, the A10P mutation reduced the distance to the hydrophobic pocket and enhanced hydrophobicity. While this change suggested potential for improved binding, it also eliminated a critical hydrogen bond mediated by R4. The detrimental impact on this key interaction likely outweighs the modest energetic gain from the reduced binding distance, supporting its classification as a non-functional or deleterious variant.

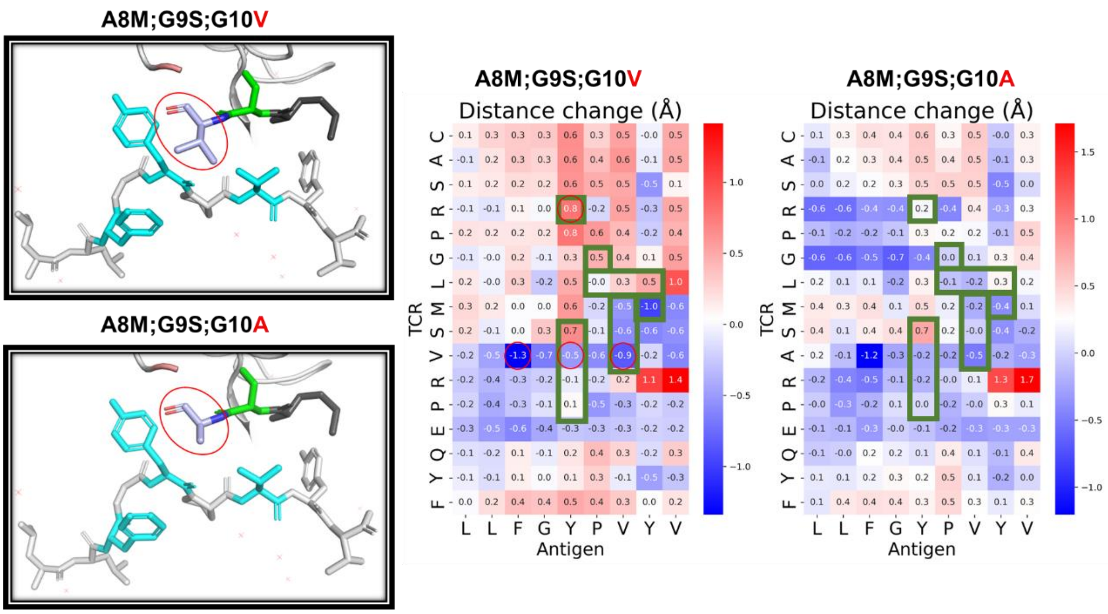


In the MSA positive control, the A10V mutation reduced the distance to the hydrophobic pocket and enhanced hydrophobicity. While this change suggested potential for improved binding, it also eliminated a critical hydrogen bond mediated by R4, supporting its classification as a non-functional or deleterious variant.

A comprehensive evaluation of random substitutions at positions 8, 9, and 10 confirmed that the original MSA sequence remains the optimal conformation for preserving functional integrity.

##### 3.2. Target Positions: 9, 10, 11

- **Ranking list**

| Rank | Mutant_ID | Mutated_Sequence | Mutation_Sites | Positions | Original_AAs | Mutated_AAs | Score |
|---|---|---|---|---|---|---|---|
| 1 | mutant_006503 | CASRPGLASWQPEQYF | G9S;G10W;R11Q | 9;10;11 | G;G;R | S;W;Q | 14.34042 |
| 2 | mutant_006446 | CASRPGLASPQPEQYF | G9S;G10P;R11Q | 9;10;11 | G;G;R | S;P;Q | 12.97077 |
| 3 | mutant_006447 | CASRPGLASPEPEQYF | G9S;G10P;R11E | 9;10;11 | G;G;R | S;P;E | 12.80281 |
| 4 | mutant_006218 | CASRPGLASRQPEQYF | G9S;G10R;R11Q | 9;10;11 | G;G;R | S;R;Q | 12.58319 |
| 5 | mutant_001088 | CASRPGLAGWQPEQYF | G10W;R11Q | 10;11 | G;R | W;Q | 12.42572 |
| 6 | mutant_006808 | CASRPGLATPEPEQYF | G9T;G10P;R11E | 9;10;11 | G;G;R | T;P;E | 12.40746 |
| 7 | mutant_006541 | CASRPGLASVQPEQYF | G9S;G10V;R11Q | 9;10;11 | G;G;R | S;V;Q | 12.38964 |
| 8 | mutant_006408 | CASRPGLASMQPEQYF | G9S;G10M;R11Q | 9;10;11 | G;G;R | S;M;Q | 12.14805 |
| 9 | mutant_002893 | CASRPGLACWQPEQYF | G9C;G10W;R11Q | 9;10;11 | G;G;R | C;W;Q | 12.00201 |
| 10 | mutant_006332 | CASRPGLASHQPEQYF | G9S;G10H;R11Q | 9;10;11 | G;G;R | S;H;Q | 11.80695 |
| 11 | mutant_005820 | CASRPGLAFVEPEQYF | G9F;G10V;R11E | 9;10;11 | G;G;R | F;V;E | 11.39525 |
| 12 | mutant_002837 | CASRPGLACPEPEQYF | G9C;G10P;R11E | 9;10;11 | G;G;R | C;P;E | 11.27354 |
| 13 | mutant_006294 | CASRPGLASQQPEQYF | G9S;G10Q;R11Q | 9;10;11 | G;G;R | S;Q;Q | 11.20895 |
| 14 | mutant_006199 | CASRPGLASAQPEQYF | G9S;G10A;R11Q | 9;10;11 | G;G;R | S;A;Q | 11.17414 |
| 15 | mutant_005591 | CASRPGLAFEQPEQYF | G9F;G10E;R11Q | 9;10;11 | G;G;R | F;E;Q | 11.12528 |
| 16 | mutant_005819 | CASRPGLAFVQPEQYF | G9F;G10V;R11Q | 9;10;11 | G;G;R | F;V;Q | 11.09242 |
| 17 | mutant_001033 | CASRPGLAGPGPEQYF | G10P;R11G | 10;11 | G;R | P;G | 11.03172 |
| 18 | mutant_005724 | CASRPGLAFPQPEQYF | G9F;G10P;R11Q | 9;10;11 | G;G;R | F;P;Q | 10.93725 |
| 19 | mutant_001031 | CASRPGLAGPQPEQYF | G10P;R11Q | 10;11 | G;R | P;Q | 10.87103 |
| 97 | mutant_006200* | CASRPGLGSAEPEQYF | G9S;G10A;R11E | 9;10;11 | G;G;R | S;A;E | 7.832404 |

***Positive control**: This mutant exhibited a significant enhancement in activity against the Tax peptide, with a calculated ΔΔG of -3.49 kcal/mol.

#### ◆ Motif Analysis

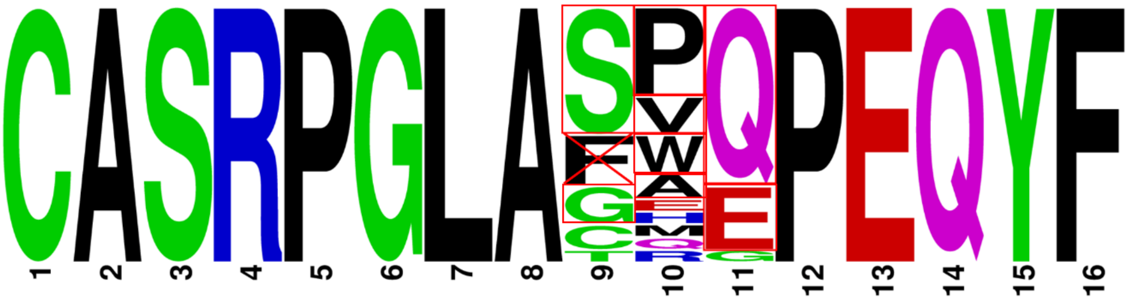

i. Position 9: Strong preference for polar residues (**S, G**)

ii. Position 10: Strong preference for polar residues (**P, V, W, A**)

iii. Position 11: Strong preference for **Q** or **E** residues.

**iv. Conserved Motifs in Top Candidates: [S/G]-[P/V/W/A]-[Q/E]**

#### ◆ Candidate Mutants Evaluation

G9S-G10P-R11Q; G9S-G10P-R11E; G9S-G10V-R11Q; G9S-G10W-R11Q; G9S-G10A-R11Q; G9S-G10A-R11E (*positive control).

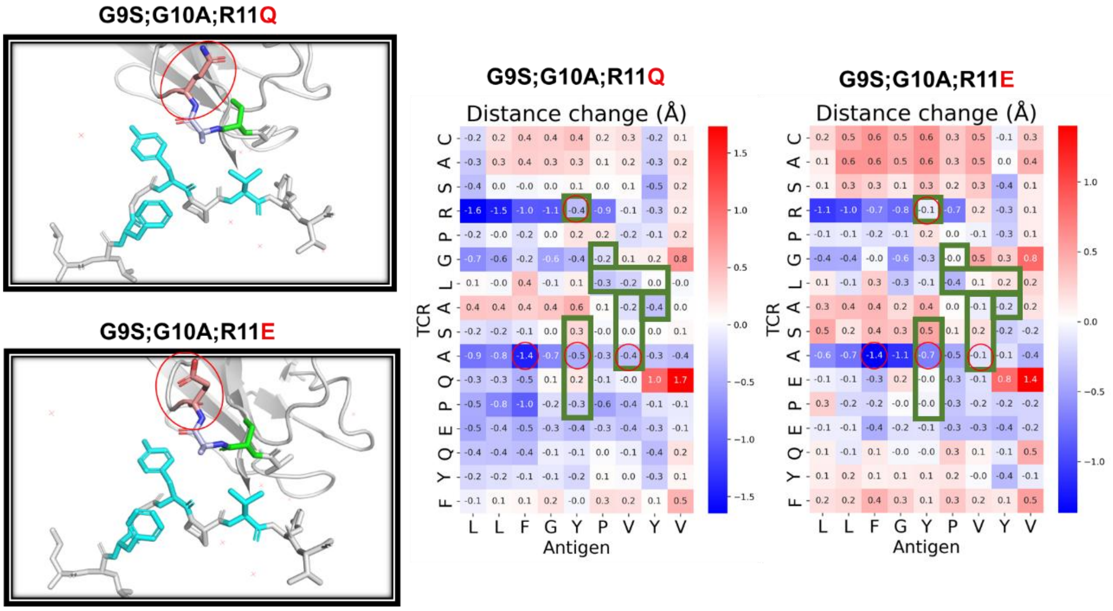


In the SAE positive control, the E11Q mutation maintained the original binding mode of A10 while significantly shortening the interaction distance with R4. This structural optimization enhances the stability of hydrogen bonding, supporting its classification as a beneficial mutation.

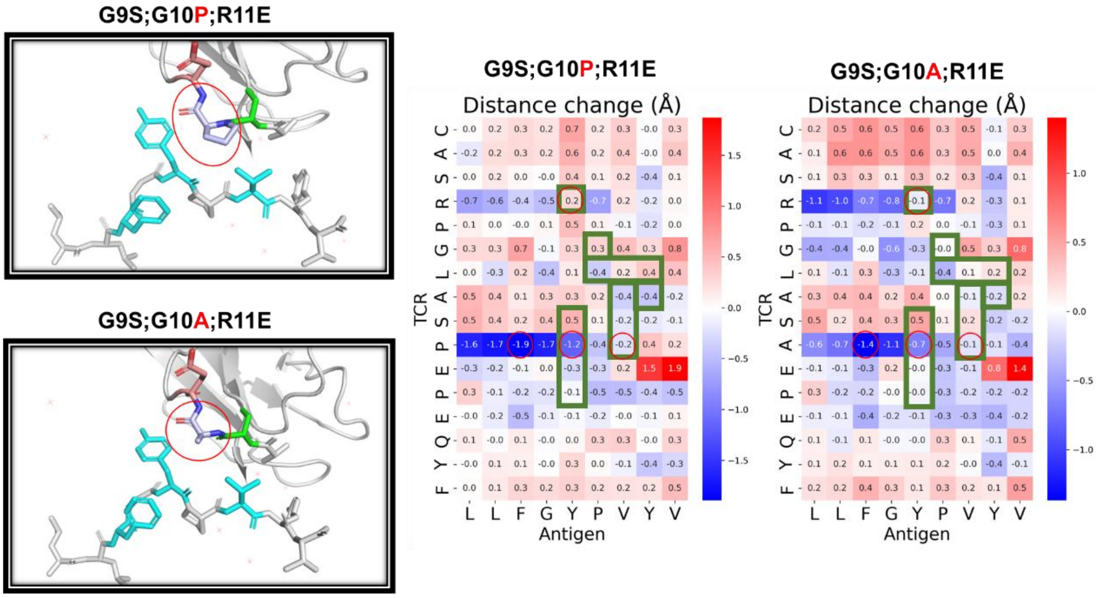


In the SAE positive control, substitution of alanine (A) at position 10 with proline (P) shortened the binding distance to the hydrophobic pocket that enhanced hydrophobic interactions, while preserving the critical hydrogen bond with arginine (R) at position 4, supporting its classification as a beneficial mutation.

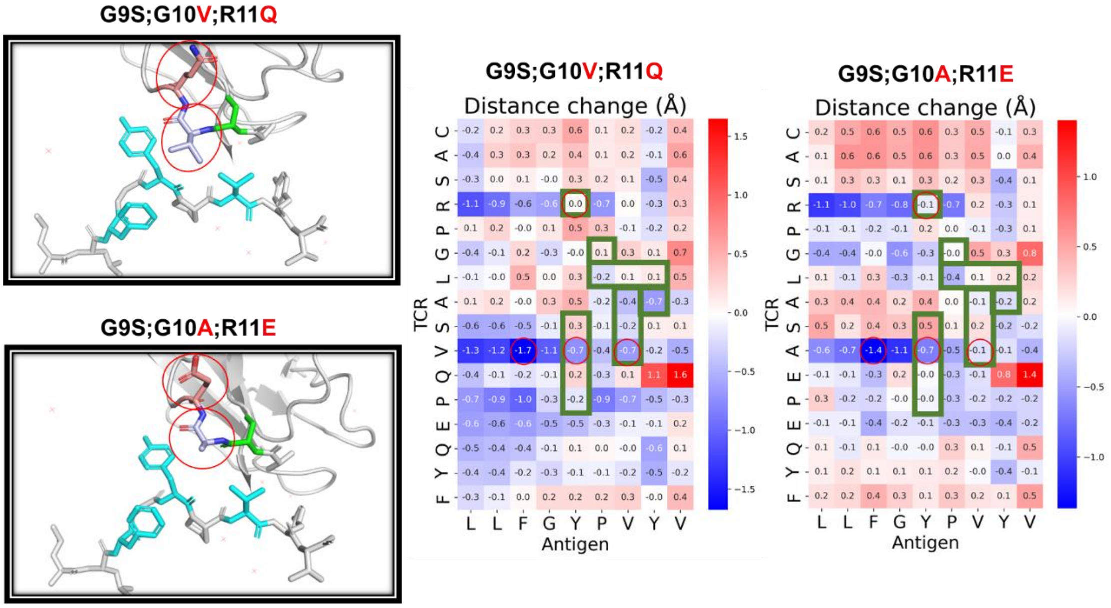


In the SAE positive control, the combined A10V and E11Q mutations resulted in reduced binding distance to the hydrophobic pocket while maintaining the critical interaction with arginine (R) at position 4, supporting its classification as a beneficial mutation.

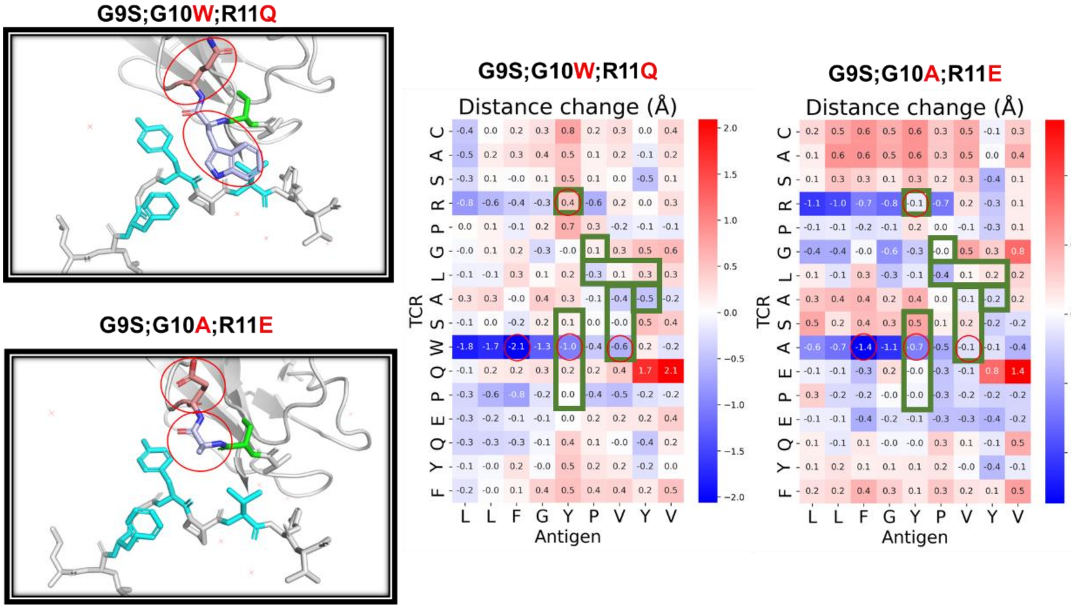


In the SAE positive control, the combined A10W and E11Q mutations reduced the binding distance to the hydrophobic pocket. However, the bulky side chain of W10 likely causes steric hindrance and disrupts the critical hydrogen bond with arginine (R) at position 4. The structural destabilization caused by this mutation outweighs the potential benefit of the closer binding proximity, supporting its classification as a non-functional or deleterious variant.

**A comprehensive evaluation of random substitutions at positions 9, 10, and 11 identifies SAQ, SPE, and SVQ as promising candidate sequences with potential performance superior to the original SAE positive control.**
